# Spectral clustering of single-cell multi-omics data on multilayer graphs

**DOI:** 10.1101/2022.01.24.477443

**Authors:** Shuyi Zhang, Jacob R. Leistico, Raymond J. Cho, Jeffrey B. Cheng, Jun S. Song

## Abstract

Single-cell sequencing technologies that simultaneously generate multimodal cellular profiles present opportunities for improved understanding of cell heterogeneity in tissues. How the multimodal information can be integrated to obtain a common cell type identification, however, poses a computational challenge. Multilayer graphs provide a natural representation of multi-omic single-cell sequencing datasets, and finding cell clusters may be understood as a multilayer graph partition problem.

We introduce two spectral algorithms on multilayer graphs, spectral clustering on multilayer graphs (SCML) and the weighted locally linear (WLL) method, to cluster cells in multi-omic single-cell sequencing datasets. We connect these algorithms through a unifying mathematical framework that represents each layer using a Hamiltonian operator and a mixture of its eigenstates to integrate the multiple graph layers, demonstrating in the process that the WLL method is a rigorous multilayer spectral graph theoretic reformulation of the popular Seurat weighted nearest neighbor (WNN) algorithm. Implementing our algorithms and applying them to a CITE-seq dataset of cord blood mononuclear cells yields results similar to the Seurat WNN analysis. Our work thus extends spectral methods to multimodal single-cell data analysis.

The code used in this study can be found at https://github.com/jssong-lab/sc-spectrum

## 1 Introduction

Graph-based data analysis and machine learning methods have been successfully used for compressed representation of data and clustering (Ng *et al*., 2001; Belkin and Niyogi, 2003; von Luxburg, 2007; Dong *et al*., 2013; McInnes *et al*., 2018b; Zhao and Song, 2018; El Gheche *et al*., 2020). The mathematical structure of graphs greatly facilitates the analysis of real-world networks such as computer, gene regulatory, protein-protein interaction, and social networks. Graph-based methods are particularly well suited for exploring high-dimensional data in single-cell genomics by succinctly capturing the results of profiling genome-wide transcripts, chromatin accessibility, or surface proteins in tens of thousands of individual cells via next-generation sequencing (NGS)-based technologies. By constructing a similarity graph, with each node representing a cell and each link between two nodes representing their similarity, information about the neighborhood of cells on the graph can be used to capture distinct cell types reflected in the clustering structure of nodes. Using pairwise similarities between data points can also help reduce the dimension of data; e.g., t-SNE (t-Distributed Stochastic Neighbor Embedding) (van der Maaten and Hinton, 2008) and UMAP (Uniform Manifold Approximation and Projection) (McInnes *et al*., 2018b) are two nonlinear dimension reduction methods based on pairwise similarities and have been extensively used for visualizing single-cell data.

Recent technological advances have enabled simultaneous generation of multi-omic data in single cells. For example, technologies like cellular indexing of transcriptomes and epitopes by sequencing (CITE-seq) (Stoeckius *et al*., 2017; Liu *et al*., 2020), as well as RNA expression and protein sequencing (REAP-seq) (Peterson *et al*., 2017) and AbSeq (Shahi *et al*., 2017), have made it possible to simultaneously perform single-cell RNA profiling and immunophenotyping (via profiling surface proteins). The rapid generation of single-cell multi-omic data has stimulated interest in developing graph-based algorithms that can integrate multimodal cellular profiles for dimensionality reduction and improved cell type identification. For example, CiteFuse (Kim *et al*., 2020) integrates the transcriptome and epitope tagging information of CITE-seq by using a similarity network fusion algorithm (Wang *et al*., 2014). The t-SNE and UMAP algorithms have been also generalized to handle single-cell multimodal data (Do and Canzar, 2021). More recently, the weighted nearest neighbor (WNN) algorithm, built on top of the popular Seurat workflow for analyzing individual assays (Stuart *et al*., 2019), utilizes a weighted summation of the resulting multiple graphs to obtain a joint-similarity WNN graph (Hao *et al*., 2021); the key idea is to perform within-modality and cross-modality predictions and to assign cell-specific modularity weights based on these predictions to construct the WNN graph.

Spectral clustering is currently one of the state-of-the-art algorithms for graph partitioning and can also facilitate dimension reduction. This algorithm does not make strong assumptions about the shape of clusters in the original point cloud and can be very fast for analyzing large sparse graphs (Ng *et al*., 2001; von Luxburg, 2007). While different variations exist, spectral clustering typically involves constructing a second-order difference operator called the graph Laplacian and using its first few lowest-frequency eigenvectors to embed and cluster the graph nodes in a low-dimensional space (Roweis and Saul, 2000; Ng *et al*., 2001; Belkin and Niyogi, 2003; Dhillon *et al*., 2007). Spectral analysis of graphs is incorporated into many algorithms and workflows. For example, the recently developed UMAP algorithm for dimension reduction (McInnes *et al*., 2018b) is built upon the same mathematical foundations as the previous Laplacian Eigenmaps (Belkin and Niyogi, 2003). CiteFuse also uses spectral clustering to partition the fused graph and has shown that it outperforms alternative methods (Kim *et al*., 2020).

A multilayer graph is a set of graphs defined on the same nodes and can simultaneously represent several multi-omic single-cell sequencing datasets. Spectral methods on a multilayer graph may then help cluster the cells by integrating information across individual graphs. One promising approach is the Spectral Clustering on Multi-Layer graphs (SCML) algorithm that finds a compromise among the low-frequency eigenspaces of individual graphs represented as points on a Grassmannian manifold (Dong *et al*., 2013). The SCML algorithm seeks a consensus subspace that does not deviate too much from any one of the individual eigenspaces on this manifold.

The main goal of this paper is to unify the SCML (Dong *et al*., 2013) and the WNN (Hao *et al*., 2021) algorithms, two seemingly different approaches to identifying communities on multilayer graphs, into a common mathematical framework. We first propose two versions of Hamiltonian on a single-layer graph, both being symmetric positive semi-definite matrices constructed from the graph Laplacian and related to spectral clustering and locally linear embedding (Roweis and Saul, 2000), respectively; we demonstrate that each Hamiltonian and an associated ensemble of eigenvectors, encoded as a density matrix, provide a suitable representation of the graph. A density matrix plays a role similar to a mixture distribution of probabilities in statistics. Intuitively, the density matrix in our formalism encodes the correlation structure of nodes in the spectral embedding space obtained from the low-frequency eigenvectors of the Hamiltonian, while the Hamiltonian itself encodes the local connectivity of nodes on the original graph. Using this formalism, we interpret the SCML algorithm and reformulate the WNN analysis as optimization problems involving a set of Hamiltonian operators and density matrices from the individual layers of a multilayer graph, thereby providing a unified theoretical foundation for these methods and explaining their similarities and differences. Applying the SCML algorithm and the reformulated WNN method to a single-cell CITE-seq dataset consisting of human cord blood mononuclear cells (CBMCs) (Stoeckius *et al*., 2017) yields results comparable to those obtained using the original WNN analysis.

## 2 Methods

### 2.1 Mathematical structure of graphs

A graph is a pair 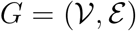, where 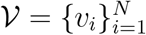 is the set of nodes (vertices), and 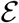 the set of links (edges) between nodes. A simple graph (without self-loops and multiple edges) can be described by an adjacency matrix **A**, with the element *A_ij_* ≥ 0 representing the strength of the link from node *v_i_* to node *v_j_*; *A_ij_* = 0 means the link does not exist. This paper considers undirected weighted graphs only and thus assumes *A_ij_* = *A_ji_*. The nodes often represent data points, and the weighted edges represent some measure of pairwise similarity between the data points.

The graph Laplacian is defined as **L** = **D – A**, where the degree matrix **D** is diagonal with entries

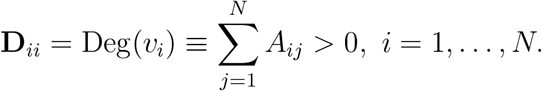

The graph Laplacian is a discrete version of the Laplace operator on manifolds (Chung, 1997; Belkin and Niyogi, 2003; von Luxburg, 2007), up to a minus sign. The set of all real-valued functions on the vertex set 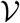 is isomorphic to 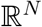. Thus, *M* functions 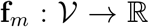, *m* = 1, …, *M*, defined on 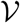 can be organized into an *N* × *M* matrix *F* = [**f**_1_, …, **f**_*m*_, …, **f**_*M*_], which may be viewed as a 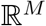-valued function or a vector field on 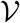. Let *F*(*v_i_*) denote the *i*-th row of this matrix. Applying the graph Laplacian **L** to this matrix gives

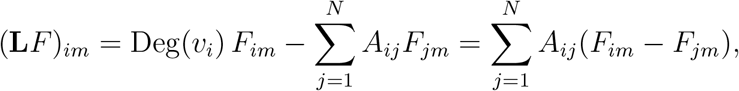

and since we assume that **A** = **A**^T^, we have

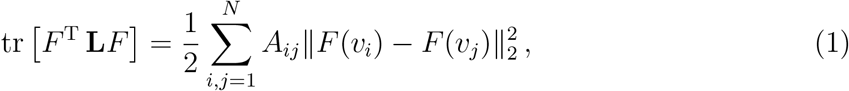

which is just the sum of squared difference of vectors at two nodes weighted by the adjacency *A_ij_*. It resembles the kinetic energy for a vector field and indicates how smoothly its components are varying on the graph.

A multilayer graph of *s*_max_ layers, 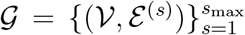, is a set of *s*_max_ graphs sharing a common node set 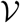. The individual graphs, 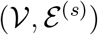, are referred to as graph layers and represent different connectivity between the nodes in 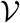, as described by the respective adjacency matrices 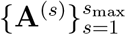 or, equivalently, the graph Laplacian operators 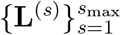. In the context of multi-omic single-cell datasets, different graph layers may encode the similarity of cells measured by different modalities. For example, from CITE-seq data, we can calculate two graph Laplacians from the RNA layer and the protein layer (Fig. 1A). The diagonal terms come from the degree matrix, and different nodes, or even the same node in different layers, may have different degrees. The off-diagonal terms arising from subtracting the adjacency matrix are non-positive. Different layers of a multilayer graph may appear quite different, as links present in one graph may be absent in others (Fig. 1A).

**Figure 1:**
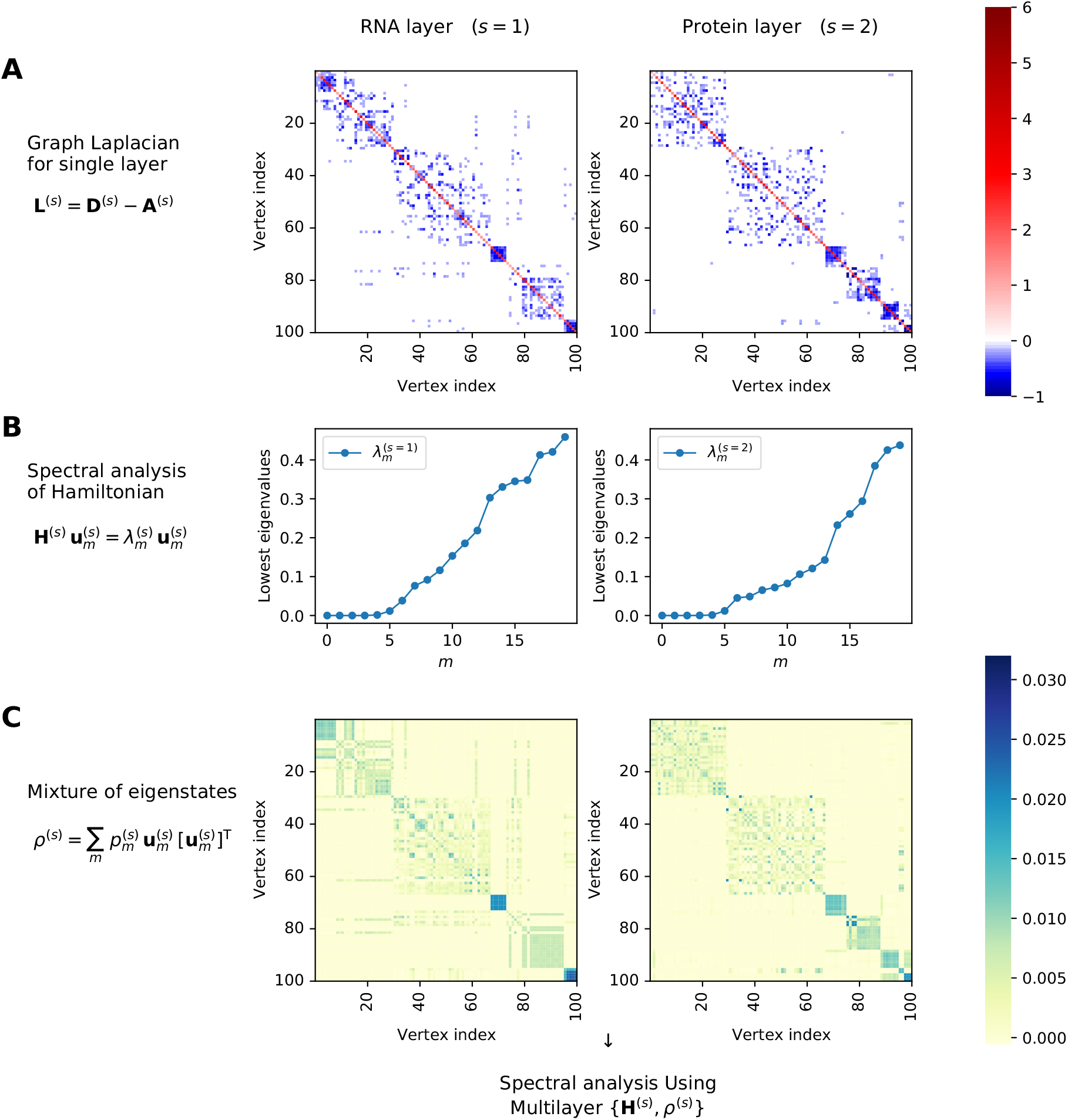
Illustration of the spectral analyses underlying our interpretation of the SCML method and reformulation of the WNN method. A subset of *N* = 100 cells were randomly selected from the CITE-seq data of human cord blood mononuclear cells. (A) Heatmap display of the graph Laplacian **L**^(*s*)^ constructed from the data in each modality. (B) The spectrum of the Hamiltonian **H**^(*s*)^ constructed from the graph Laplacian of each layer. The locally linear construction from Equation (4) with **W** = **I** is shown. (C) Heatmap display of a density matrix representing a mixture of Hamiltonian eigenstates for each layer. The thermal mixture from Equation (8) is shown.

One simple method of integrating the layers would be to take the arithmetic mean of the graph Laplacians 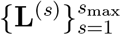. However, this approach would amount to averaging the pairwise similarities without considering the higher-order structure of individual graphs. Other approaches include making the graph Laplacians commute approximately (Bronstein *et al*., 2013), or equivalently, jointly diagonalizing the graph Laplacians approximately (Eynard *et al*., 2015), as well as finding the geometric mean of the graph Laplacians (El Gheche *et al*., 2020).

Motivated by the SCML algorithm (Dong *et al*., 2013), this paper proposes a spectral method for merging multiple graph layers, each of which is represented by a Hamiltonian operator and a mixture of its eigenstates. The following section introduces two ways of constructing the Hamiltonian for a single-layer graph, and then justifies using the density matrix to capture the clustering structure of nodes on the graph.

### 2.2 Spectral analysis of graphs and mixture of eigenstates

As seen in Equation (1), the graph Laplacian behaves like a discrete version of the Laplace operator on functions defined on the set of nodes. Starting from the graph Laplacian, we construct two different Hamiltonian operators corresponding to the different optimization problems associated with the SCML and Seurat WNN algorithms.

The first Hamiltonian is closely related to spectral clustering and the problem of minimizing weighted cuts on graphs (von Luxburg, 2007; Meilă and Pentney, 2007; Dhillon *et al*., 2007):

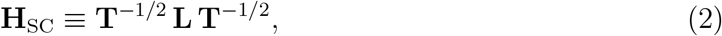

where 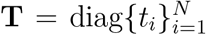, *t_i_* > 0, ∀*i*, is a diagonal matrix of positive values describing some notion of “importance” of the corresponding *N* nodes. Without further information about a graph, we will choose **T** to be the degree matrix:

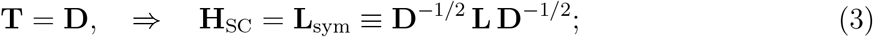

calculating a low-frequency eigenspace (a vector space spanned by the first several eigenvectors associated with the lowest eigenvalues) of this symmetrically normalized graph Laplacian provides an approximate solution minimizing the normalized cuts on the graph (Shi and Malik, 2000; Yu and Shi, 2003). The normalized cuts criterion aims to partition a graph by minimizing the weights of intercluster links, normalized by the weights of intracluster links (von Luxburg, 2007).

The second Hamiltonian motivated by the Locally Linear Embedding (LLE) algorithm (Roweis and Saul, 2000) is:

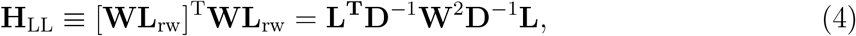

where **L**_rw_ ≡ **D**^−1^**L** is the random walk graph Laplacian (Supplementary Methods), and 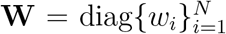, *w_i_* ∈ [0,1], ∀*i*, is another diagonal matrix of node-specific weights. The random walk graph Laplacian computes the difference between a function’s true value at a node and a weighted average of its values at immediate neighbors, as follows

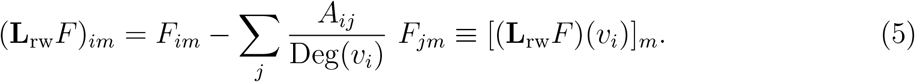

The Locally Linear Hamiltonian thus yields the quadratic form

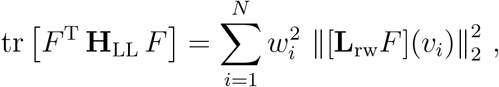

which sums the squared neighborhood prediction error at each node *v_i_* weighted by 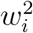. In the absence of further information, a natural choice for **W** is **W** = **I**, yielding **H**_LL_ = [**L**_rw_]^T^**L**_rw_, as proposed by the original LLE algorithm (Roweis and Saul, 2000). However, when there is more than one graph layer, these node-specific weights for each layer can be learned from the data and play a key role in understanding the WNN analysis.

For either choice of the Hamiltonian operator **H**, suppose the eigenvalues are ordered as 0 ≤ λ_0_ ≤ λ_1_ ≤ ⋯ ≤ λ_*N*–1_, with corresponding orthonormal eigenvectors **u**_0_, **u**_1_, …, **u**_*N*–1_. The matrix

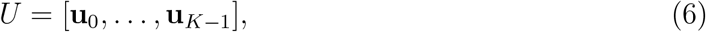

forms the solution to the trace minimization problem

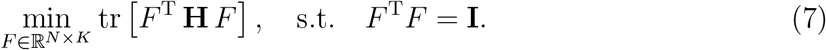

For the Spectral Clustering Hamiltonian **H**_SC_, the rows of *U* provide the spectral embedding (Belkin and Niyogi, 2003) and facilitate the spectral clustering (Ng *et al*., 2001) of the nodes. For the Locally Linear Hamiltonian **H**_LL_, the rows of *U* offer the neighborhood-preserving mapping minimizing the reconstruction errors of the embedded coordinates (Roweis and Saul, 2000). The spectrum of **H**_LL_, with **W** = **I**, for the CITE-seq data is shown in Fig. 1B.

A density matrix is a matrix that represents an ensemble of quantum states, similar to a mixture distribution of probabilities. In this formalism, a pure state corresponding to a vector **v** is represented by its outer product with itself, **vv**^T^, while a mixed state is represented by a sum of pure states weighted by their mixing probabilities. We can thus encode a mixture of Hamiltonian eigenstates (pure states corresponding to Hamiltonian eigenvectors **u**_*m*_, *m* = 0, …, *N* – 1) as the density matrix

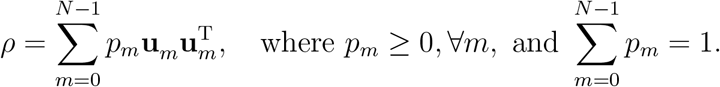

In particular, the density matrix *ρ* = *UU*^T^/*K*, recapitulating the low-frequency eigenspace representation in Equation (6), is the solution to the following trace minimization problem resembling Equation (7):

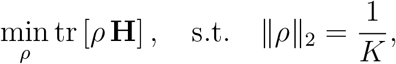

where ||*ρ*||_2_ = max*_m_*{*p_m_*} is the spectral norm of *ρ*, and 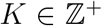 is fixed. That is, a mixed state that minimizes the expected energy under the spectral norm constraint has a uniform mixture of the lowest *K* eigenstates (Supplementary Methods).

Alternatively, we can choose a probability distribution arising from the thermal equilibrium distribution in statistical physics, or the softmax function in machine learning:

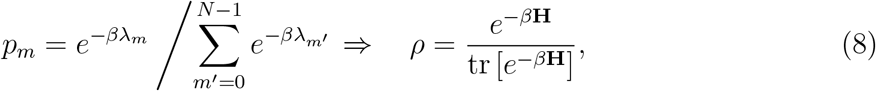

where *β* = 1/*T* is the inverse temperature. The thermal equilibrium density matrix is the solution to a trace minimization problem over density matrices with a different constraint,

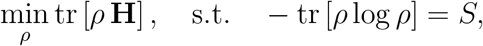

where *S* > 0 is fixed entropy, and the Lagrange multiplier *β*^−1^ satisfies the relation *S* = *β* tr [*ρ***H**]+log tr [*e*^−*β***H**^] (Supplementary Methods). Equivalently, this distribution maximizes entropy given fixed ensemble average of energy and provides the most unbiased probabilistic description of biological data (Finnegan and Song, 2017). We recommend choosing the value 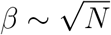; see Supplementary Methods and Supplementary Figures 1, 2, 3 for discussions on potential effects of varying *β* on clustering results.

For the RNA and protein layers of the CITE-seq data, Fig. 1C shows their thermal equilibrium density matrices using the respective **H**_LL_. In the Results sections 3.1 and 3.2, we will make specific choices of {**H**^(*s*)^, *ρ*(*s*)} for each layer to interpret the SCML algorithm and reformulate the WNN analysis in a common mathematical framework.

To obtain a feature map from the density matrix 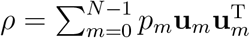, we define

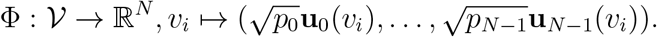

Using this notation, the density matrix can be expressed as *ρ* = ΦΦ^T^, encoding the correlation structure of nodes in the embedding space.

### 2.3 CBMC CITE-seq data preprocessing

The scRNA-seq unique molecular identifier (UMI) count matrix was obtained from the Gene Expression Omnibus (GEO): GSE100866, `GSE100866_CBMC_8K_13AB_10X-RNA_umi.csv.gz`. Human genes were identified as those that began with “HUMAN_”, and human cells were identified as those with greater than 90% of UMI tags aligned to human genes, resulting in a filtered UMI matrix, 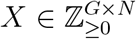, of *G* = 20400 human genes and *N* = 8005 human cells. We normalized the filtered count matrix by library size to remove the effect of variable transcript counts, and applied a square root transform to reduce the effect of extreme values, as follows:

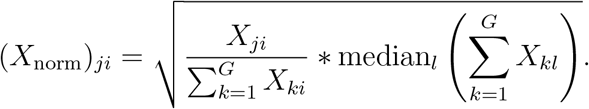

Finally, we performed principal component analysis (PCA) to denoise the high dimensional normalized expression data by projecting each cell onto the top 30 principal components. PCA was performed on the transpose of *X*_norm_ using sklearn.decomposition.PCA with options (n_components = 30, random_state = 12345678).

The antibody-derived tag (ADT) UMI matrix was also obtained from the GEO, `GSE100866_CBMC_8K_13AB_10X-ADT_umi.csv.gz`. We filtered this matrix to contain only the human cells identified from the scRNA-seq UMI count matrix, resulting in a count matrix 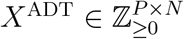, where *P* = 13 is the number of antibodies profiled and *N* = 8005 is the number of human cells. The cell ADT profiles were normalized using a centered log ratio (CLR) transform with a pseudo-count of 1 (Stoeckius *et al*., 2017):

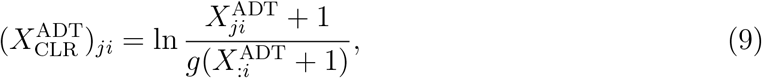

where 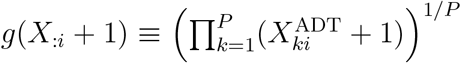 is the geometric mean of pseudo count adjusted ADT UMIs for cell *i*.

### 2.4 Graph construction

We constructed a cell-cell similarity graph using a method motivated by (van Dijk *et al*., 2018). First, a *k* nearest neighbor (*k*-NN) distance graph was constructed for each modality using the Euclidean distance between the cell normalized profiles (PCA for RNA and CLR transform for ADT). The nearest neighbor distance graph was obtained using sklearn.neighbors.NearestNeigherbors with options (m_neighbors = *k*, metric = “minkowski”, p = 2). A similarity graph was obtained from the *k*-NN distance graph using an adaptive Gaussian radial basis function. To construct the similarity graph, let *D* denote the cell-cell pairwise distance matrix and *k*NN_*i*_(*j*) denote the *j*th neighbor of cell *i* for a given modality (RNA or ADT). The nonzero entries of the corresponding similarity graph *B* were obtained as

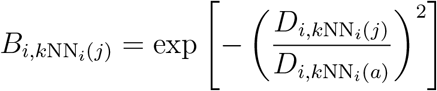

where *a* ∈ [1, *N*] is a hyperparameter that specified the adaptive kernel width. For all datasets, we used the values *k* = 30 and *a* = 10, as recommended by (van Dijk *et al*., 2018), except in Fig. 1 where *N* = 100 cells were subsampled and in Supplementary Figures 1-3 where *N* = 100 data points were simulated, in which case *k* = 5 and *a* = 5 were used. All other entries in the matrix were set to zero. A symmetric graph adjacency matrix **A** was obtained by symmetrizing the *k*-NN similarity matrix as 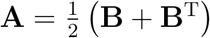.

### 2.5 Spectral clustering implementation

Spectral decomposition and clustering of a cell similarity graph was performed on the corresponding symmetric graph Laplacian defined in in Equation (3). The spectral decomposition of the symmetric graph Laplacian was performed using scipy.sparse.linalg.eigsh. For clustering into *K* clusters, *K* orthonormal eigenvectors with the smallest eigenvalues were stacked as columns in 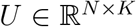. The spectral embedding vectors for each cell were *ℓ*_2_ normalized for the purpose of clustering (Ng *et al*., 2001; von Luxburg, 2007)

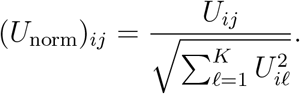

Clustering was performed using sklearn.cluster.KMeans with options (n_cluster = *K*, max_iter = 1000, n_init = 100, random_state = 12345678).

### 2.6 Spectral clustering on multilayer graphs (SCML)

The SCML method was implemented for CITE-seq data by first constructing the symmetric graph Laplacians for the RNA and ADT profiles 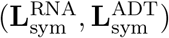. The *K* eigenvectors with smallest corresponding eigenvalues were then obtained for each of the symmetric graph Laplacians, yielding two unnormalized spectral embedding matrices (*U*^RNA^, *U*^ADT^). The modified SCML graph Laplacian (Dong *et al*., 2013) with hyperparameter 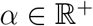 was then constructed:

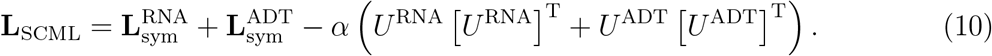

The spectral decomposition of the modified Laplacian was performed using Pytorch (torch.linalg.eigh) after first converting the modified Laplacian to a Pytorch Tensor object. Cell clustering was performed with the *K* eigenvectors obtained from this decomposition using the same normalization and clustering described above.

### 2.7 UMAP visualization

UMAP (McInnes *et al*., 2018b) was used to visualize a 2-dimensional embedding of the ℓ_2_-normalized SCML embedding vectors (Methods 2.5). The 2-dimensional UMAP projections were constructed using the umap-learn UMAP function (McInnes *et al*., 2018a) with options (random_state = 12345678, n_neighbors = 30, set_op_mix_ratio = 1, local_connectivity = 1, min_dist = 0.3, metric = ”cosine”).

### 2.8 Silhouette scores

The full pairwise distance matrices for the RNA PCA and ADT CLR profiles were calculated using scipy.spatial.distance.pdist with options (metric = ”euclidean”) followed by scipy.spatial.distance.squareform to format the distances as a matrix. The silhouette scores for the RNA and ADT distance matrices were then obtained with sklearn.metrics.silhouette_score with option (metric = “precomputed”).

### 2.9 Seurat analysis of the CBMC dataset

Using the same criteria as in Methods 2.3, we selected the cells with more than 90% RNA reads aligned to the human genome, and obtained the RNA and ADT count data for a total of *N* = 8005 human cells, removing the RNA count from the mouse genes (using the ”CollapseSpeciesExpressionMatrix” function with ncontrols = 0). We also removed three insignificant ADTs (CCR5, CCR7, CD10), as previously suggested (Stuart *et al*., 2019).

We then followed the standard Seurat workflow for CITE-seq data. Analysis of the RNA data involved: normalizing the count data (NormalizeData) using with the default normalization.method = ”LogNormalize”, scale.factor = 10000; finding the 2000 most variable features; scaling those selected features; and, running PCA. Analysis of the ADT data involved: normalizing the count data (NormalizeData) using normalization.method = ”CLR”, margin = 2; scaling all features; and, running PCA.

Lastly, we applied the WNN method, which first constructed nearest neighbor graphs on the first 30 PCs of the RNA data and the first 7 PCs of the ADT data, then computed the cell-specific modularity weights for each modality, and built a WNN graph, all incorporated in the ”FindMultiModalNeighbors” function of Seurat v4 (Hao *et al*., 2021). The WNN graph was used for dimension reduction and clustering: a UMAP embedding in two dimensions, and a modularity-based clustering with resolution 0.2. The choice of resolution 0.2 was made to reproduce the WNN clustering result of 12 human cell type clusters reported in (Hao *et al*., 2021). The clusters are then manually labeled based on their marker genes and proteins (using the ”FindAllMarkers” function).

## 3 Results

### 3.1 SCML penalizes the Hilbert-Schmidt distance between density matrices

In the SCML algorithm for a multilayered graph with *N* vertices (Dong *et al*., 2013), the *K*-dimensional lowest-frequency eigenspace 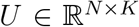 of each normalized graph Laplacian is represented as a point on the Grassmann manifold 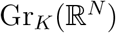 of all *K*-dimensional subspaces in 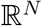. The squared chordal distance between two subspaces *U* and *U*′ on the manifold is

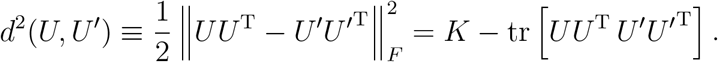

Given the individual eigenspaces *U*^(*s*)^ of *s*_max_ layers, the SCML algorithm searches for a consensus subspace *U* that minimizes the joint spectral energy (Equation (7)) of all layers while also minimizing the sum of squared distance to the eigenspace *U*^(*s*)^ of each layer, by adding the term

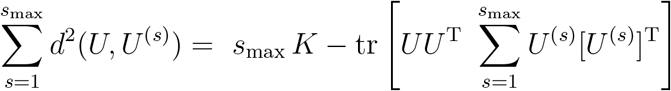

to the loss function, resulting in a new trace minimization problem with the Hamiltonian

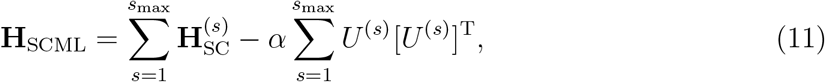

where 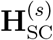 is the Spectral Clustering Hamiltonian (normalized graph Laplacian) of each layer (Equation (2)), and 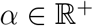 is a hyperparameter balancing the two terms in the minimization

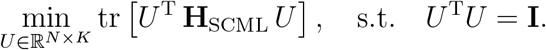

For CITE-seq data, we have *s*_max_ = 2 (*s* = 1 for RNA, *s* = 2 for ADT), and the SCML Hamiltonian **H**_SCML_ reproduces Equation (10) when the symmetrically normalized graph Laplacian is chosen for each 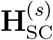.

Note that for two density matrices 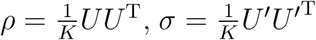 that are uniform mixtures of *K* orthonormal states, their Hilbert-Schmidt distance is equivalent to the squared distance *d*^2^(*U, U*′) on the Grassmannian manifold:

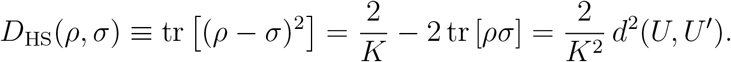

Using the uniform-mixture density matrices *ρ*^(*s*)^ associated with the *K*-dimensional lowest-frequency eigenspace of each 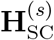, the SCML Hamiltonian becomes

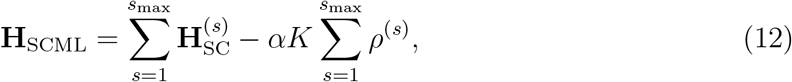

and the SCML algorithm amounts to finding a density matrix *ρ* that solves

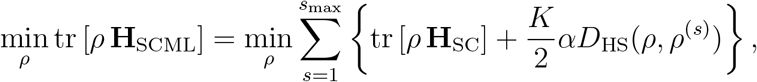

where *ρ* is again constrained to satisfy ||*ρ*||_2_ = 1/*K*. As in **H**_SC_ = **T**^−1/2^(**D** – **A**)**T**^−1/2^ where the off-diagonal contains the normalized pairwise adjacency, the magnitude of the off-diagonal terms in **H**_SCML_ (Equation (12)) also represents the modified affinity between nodes,

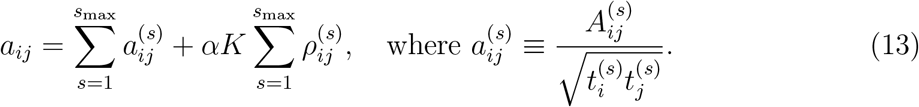

We can thus interpret the SCML algorithm as computing the modified affinity *a_ij_* between nodes on a multilayer graph, by introducing additive corrections 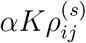 to the original affinity 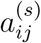. The term 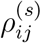, known as the spectral kernel, computes the covariance of nodes in the spectral embedding space of each layer *s*.

In next section, we will use the same language of Hamiltonian operators and density matrices to reformulate the WNN method as computing a different form of affinity between nodes on a multilayer graph.

### 3.2 Multilayer spectral graph theory provides a mathematical foundation for the WNN method

The WNN analysis aims to compute the weighted affinity between graph nodes based on their affinity in each modality (Hao *et al*., 2021), as follows:

1. For each modality, compute a similarity graph with an adjacency matrix **A**^(*s*)^ and a low-frequency representation Φ^(*s*)^ of the nodes.
2. Calculate the nearest neighbor prediction error for layers *s*_1_ and *s*_2_:

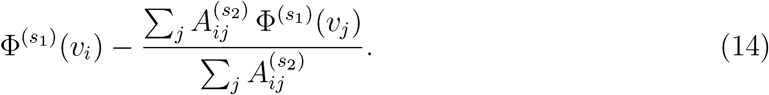 If *s*_1_ = *s*_2_, it is the within-modality nearest neighbor prediction error; otherwise, it is the cross-modality prediction error. A key observation used in our work is that we can rewrite the above prediction error as 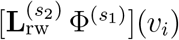, as in Equation (5).
3. From the within- and cross-modality nearest neighbor prediction errors, learn a set of modality weights 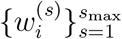 for each node *v_i_*, satisfying

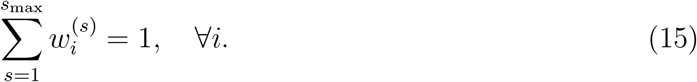
4. Calculate the weighted affinity from *v_i_* to *v_j_*, using the learned node-specific modality weights for *v_i_*,

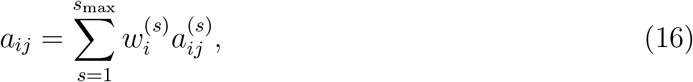

where 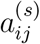 is some notion of affinity from *v_i_* to *v_j_* on graph layer *s*. Thus, the WNN approach introduces a multiplicative correction to the original affinity in each layer. Finally, a WNN graph is constructed from the weighted affinity matrix and used for downstream analysis, such as dimension reduction and clustering.

We see that in Step (2) of the WNN analysis, the random walk graph Laplacian naturally arises, suggesting that multilayer spectral graph theory may provide a rigorous mathematical foundation for the WNN method. In our ensuing interpretation of the algorithm, we make choices that utilize spectral techniques and deviate slightly from the Seurat v4 implementation (Hao *et al*., 2021). For example, the original method uses the eigenvectors from the principal component analysis (PCA) of each modality data as the representation Φ^(*s*)^ of the layer; here, we instead use a mixture of the eigenstates of a Hamiltonian operator constructed from the random walk graph Laplacian (Methods). We also formulate an optimization problem to solve for the node-specific modality weights. Despite these differences, our implementation below provides results very similar to those from the original WNN analysis.

For each modality, we first construct a similarity graph with an adjacency matrix **A**^(*s*)^ (Methods) and calculate the random walk graph Laplacian 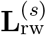. Without any prior knowledge about the graphs, we first use the identity weight matrix in the Locally Linear Hamiltonian (Equation (4)) and the thermal mixture of its eigenstates to initialize the representation of each layer:

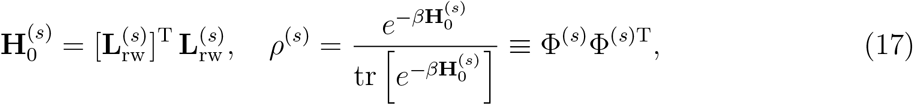

where the feature map Φ^(*s*)^ from the density matrix *ρ*^(*s*)^ embeds the set of *N* nodes in layer s into 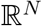 (Methods), similar to Step (1) above.

To learn the node-specific modality weights, we next modify the Locally Linear Hamiltonian to include a diagonal weight matrix, as

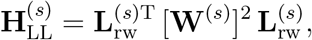

where 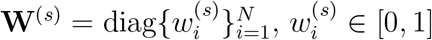, ∀*i*, *s*, with the diagonal elements satisfying Equation (15) as in Step (3) of WNN. Note that, by construction, 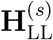 computes the weighted sum of squared within- or cross-modality prediction errors for the above feature maps (Equation (17)),

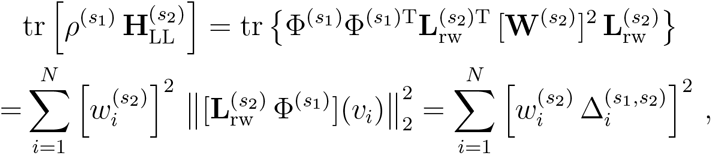

where 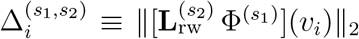. The lower the value of 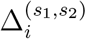, the better we can use the neighboring nodes of *v_i_* in layer *s*_2_ to predict its function value at *v_i_* in layer *s*_1_. Thus, the difference between cross- and within-modality prediction errors, weighted by 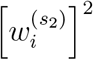, is

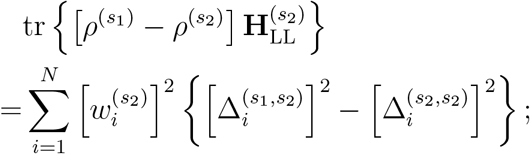

and, adding a similar term after switching *s*_1_ and *s*_2_ yields

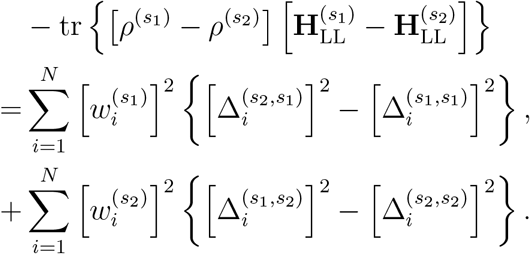

To implement Steps (2) and (3) of the WNN analysis, we propose to learn the node-specific modality weights by minimizing this difference. When *s*_max_ = 2, we thus need to solve (Supplementary Methods)

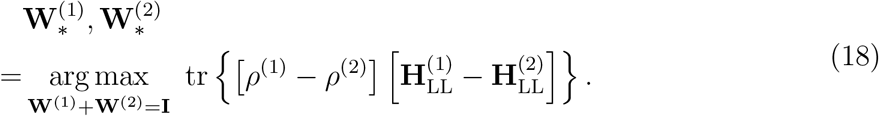

This scheme can be generalized to the case of more than two layers by iterating over each pair of layers

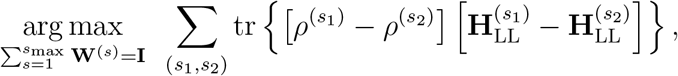

while keeping the sum of node-specific weights to be one (Supplementary Methods; Supplementary Figures 1 and 2).

Lastly, taking the affinity in Equation (16) to be 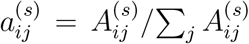, the final WNN multilayer graph can be described by the following consensus random walk graph Laplacian,

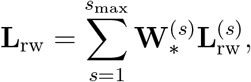

and the final Weighted Locally Linear Hamiltonian for the combined similarity graph using this consensus **L**_rw_ is

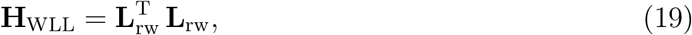

which can then be used to solve for a *K*-dimensional feature map Φ for further downstream analysis (Methods).

To compare our reformulation with the original Seurat WNN method, we applied both approaches to the human CBMC CITE-seq dataset (Stoeckius *et al*., 2017). This dataset included 5% spiked-in mouse cells, which we removed from the analysis to reduce the influence of highly distinct but biologically uninteresting cell types (Fig. 2A, Methods). The Seurat WNN clustering was performed using resolution of 0.2 (Methods), yielding 12 cell types consistent with the previously reported results (Hao *et al*., 2021). For comparison, we fixed the number of clusters to be 12 for the reformulated WNN algorithm (Fig. 2B). While our implementation of the WNN idea deviated from the Seurat software package in technical details, the clustering results were nevertheless very similar. We also showed the cell-specific weights of the RNA modality in Fig. 2C. As in the original WNN results (Hao *et al*., 2021), if the surface markers of a certain cell type were not measured in the ADT assay, then their RNA weights tended to be larger; conversely, the presence of specific surface marker measurements made the cells’ RNA weights smaller than the ADT weights.

**Figure 2:**
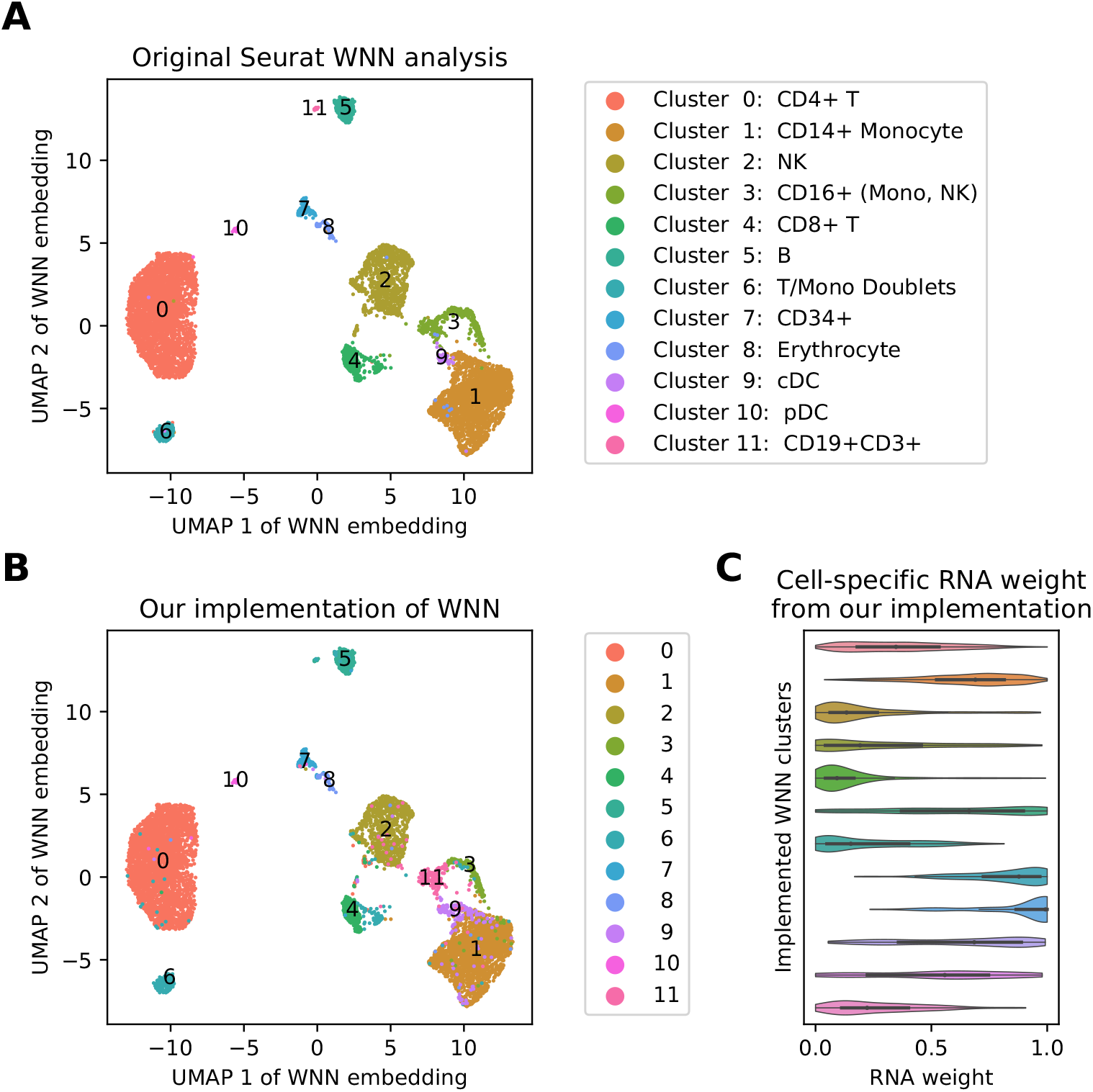
Comparison of our implementation of the WNN method with the original algorithm in Seurat v4. (A) UMAP view of the original Seurat WNN result, and a modularity-based clustering with resolution 0.2, resulting in *K* = 12 clusters, which are then manually labeled based on the marker genes and proteins. (B) The same UMAP view of the weighted locally linear (WLL) similarity graph generated by our implementation of the WNN method. Using a feature map corresponding to the *K* = 12 lowest Hamiltonian eigenstates, spectral clustering with *K*-means yields similar clustering as in (A). (C) Cell-specific modality weights obtained from our implementation show different distribution in different cell types identified in (B).

### 3.3 WNN and SCML methods yield comparable results

We compared the Seurat WNN method with graph spectral clustering approaches using the CBMC CITE-seq dataset (Stoeckius *et al*., 2017) (Methods). We partitioned the cells into 12 clusters using each of RNA spectral, ADT spectral, and SCML clustering algorithms to match the number of clusters obtained with the WNN algorithm (Methods, Supplementary Methods, Supplementary Figures 4 and 5). The clustering results for each of these four algorithms were projected onto the two-dimensional UMAP obtained from the SCML embedding vectors (Methods, Fig. 3). A discussion of how the hyperparameter *α* was chosen can be found in Supplementary Methods.

**Figure 3:**
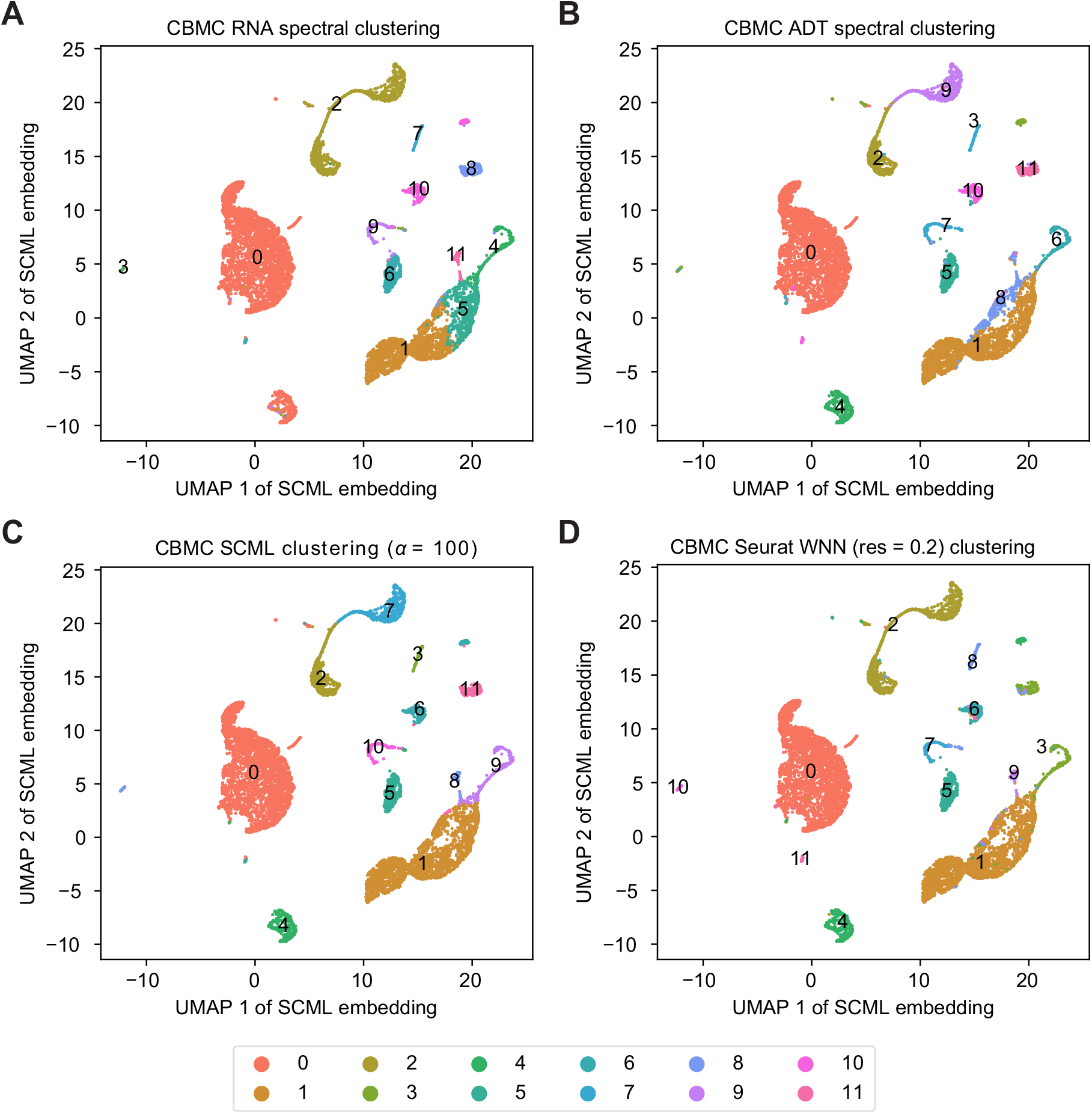
Comparison of spectral and weighted nearest neighbor clustering algorithms on the CBMC dataset. (**A**) Spectral clustering using only RNA data. (**B**) Spectral clustering using only ADT data. (**C**) SCML clustering algorithm with hyperparameter *α* = 100. (**D**) Seurat WNN algorithm with clustering hyperparameter resolution = 0.2.

We calculated the silhouette scores for both RNA and ADT pairwise distance matrices to quantify the separation of the clusters obtained using each of the four methods (Methods, Table 1). The silhouette scores demonstrated that both SCML and Seurat WNN approaches achieved a compromise between the clustering results based on either RNA or ADT alone. That is, compared to the ADT-based spectral clustering that showed a poor silhouette score for the RNA distance matrix, both integrative approaches noticeably improved the score for the RNA distance, while still maintaining a high score for the ADT distance matrix (Table 1). Similarly, compared to the RNA-based spectral clustering, both integrative approaches improved the score for the ADT distance, although the score for the RNA distance matrix decreased slightly, because the CD4+ and CD8+ T cell clusters were correctly separated by the WNN and SCML methods, as further discussed below (Table 1, Supplementary Table 1).

**Table 1:**
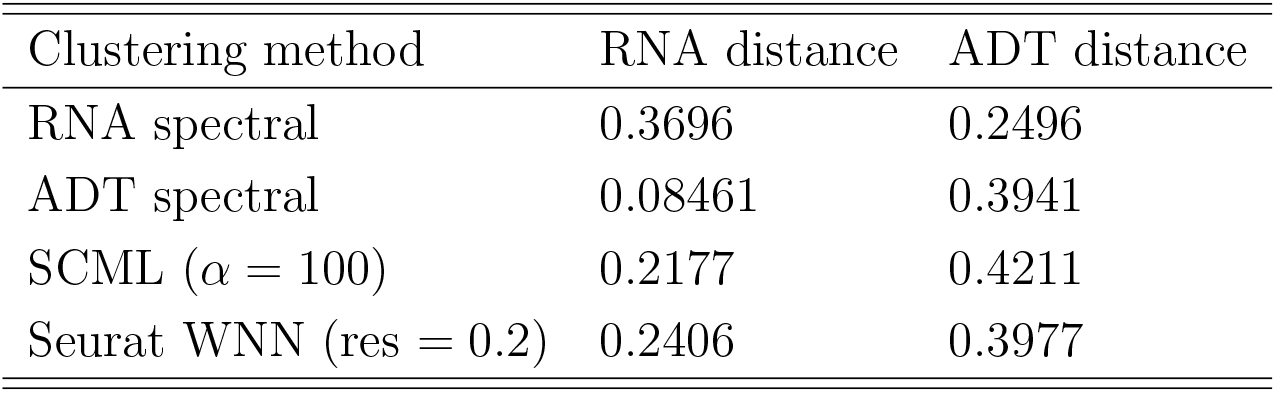
CBMC mean silhouette scores for different clustering methods and distance measures

This comparison suggested that both integrative approaches produced a clustering that compromised between the two modalities and thus likely yielded a better separation of true biological cell types, which might have been blurred by experimental biases and artifacts such as the scRNA-seq dropout effects and the limited number of profiled ADT surface proteins. For example, the RNA-based spectral clustering grouped the clusters labeled by the WNN method as CD4+ helper and CD8+ cytotoxic T cells (Seurat WNN clusters 0 and 4, respectively) (Fig. 2, Fig. 3) which were clearly separated in the ADT space. By contrast, both Seurat WNN and SCML algorithms correctly partitioned these cell types. Merging the CD4+ and CD8+ T cell clusters together improved the RNA silhouette scores, but worsened the ADT silhouette scores for both integrative methods (Supplementary Table 1), demonstrating that the ideal clustering for one modality may not correspond to that for the other and that a compromise is needed to identify biologically distinct cell types.

Comparing the clustering results obtained using the SCML algorithm (with *α* = 100) to those obtained using Seurat’s WNN algorithm (with resolution = 0.2) showed that some of the major differences involved the partitioning of natural killer (NK) cells (Seurat cluster 2) and CD16+ cells (Seurat cluster 3) (Fig. 2, Fig. 3). Each of these two cell types were further partitioned into two clusters by the SCML algorithm. The NK cells were also partitioned into two clusters by the ADT spectral clustering, but not the RNA spectral clustering, suggesting that the partitioning resulted primarily from differences in the ADT profile of cell subpopulations. Comparing ADT markers between the two partitions of the NK cells (SCML clusters 2 and 7) showed that cells in SCML cluster 7 exhibited higher levels of CD8 compared to cluster 2 (Supplementary Figure 6). A separation of the NK cells into CD8+ and CD8-subtypes in this dataset was previously described by Hao *et al*., 2021, and we also observed this separation when using a higher resolution for the WNN clustering (Supplementary Figure 7), demonstrating that the sub-partitioning of the NK cells by the SCML algorithm was actually biologically meaningful. The CD16+ cluster (Seurat cluster 3) was also partitioned into two clusters, 9 and 11, by the SCML algorithm; SCML cluster 11 exhibited a larger ADT library size than SCML cluster 9 (median total ADT UMI counts of 18008 and 2735, respectively) and elevation of CD8 and CD45RA in the CLR transformed profiles. The CD16+ cells were also partitioned into two clusters by both RNA-based and ADT-based spectral clustering, suggesting that these differences were present in both transcriptional and protein-level profiles. As was the case for the NK cells, the Seurat WNN algorithm also partitioned the CD16+ cells into two clusters, similar to those obtained by the SCML algorithm, when a larger resolution value was used (Supplementary Figure 7).

In addition to the CBMC dataset, we also compared the WNN method with spectral clustering based on RNA, ADT, and the SCML algorithm on the human bone marrow mononuclear cells (BMNC) CITE-seq data set (Stuart *et al*., 2019), clustered into 40 clusters (Supplementary Methods). Given the large number of clusters, partially overlapping colors made the results for this data set difficult to compare via simple visual inspection (Supplementary Figure 8). To compare the clustering results in a quantitative way, we thus used the normalized mutual information (NMI) score to measure the pairwise agreement between clustering methods (Supplementary Methods; Supplementary Table 2). Both Seurat WNN and SCML clustering results showed improved agreement with individual RNA and ADT modalities, and the NMI score between the Seurat WNN and SCML algorithms was the highest among all pairs of methods compared. Furthermore, the correlation between true protein ADT features and predicted profiles obtained by averaging over nearest neighbor profiles, as done in Hao *et al*. (2021), was similar for the WNN and SCML methods (Supplementary Methods; Supplementary Figure 9).

## 4 Discussion and Conclusion

In this paper, we have reformulated two algorithms, SCML and Seurat WNN, for clustering single-cell multi-omic datasets in a unified framework of multilayer spectral graph analysis. We have shown that these algorithms can be expressed as optimization problems involving different Hermitian operators acting on functions defined on the graph nodes. Our results highlight the connection between these seemingly distinct algorithms and provide a mathematical framework for understanding these multi-modal clustering algorithms. Our reformulation of the WNN method establishes a connection between this popular tool and spectral signal processing on graphs, paving the way for the future development of more advanced mathematical approaches, e.g. graph wavelet-based (Hammond *et al*., 2011) multiresolution detection of clusters on multilayer graphs. We also show that our reformulation can be easily generalized to several layers and provide an analytic formula for the weights derived from solving an optimization problem (Supplementary Methods; Supplementary Figures 1 and 2).

Both SCML and WNN algorithms modify the affinity matrix in individual layers, but they differ in motivation and implementation details. As seen in Equation (13), SCML introduces layer-specific additive corrections to the affinity matrix in each layer, where the corrections arise from the kernel evaluation of nodes in the layer-specific spectral embedding space that captures low-frequency macroscopic grouping of nodes into clusters. These corrections thus reflect the clustering preferences of individual layers, and their computation does not require coupling information across different layers. Coupling of information across layers occurs when a consensus spectral embedding space is computed to minimize the expectation value of the joint sum of the modified Hamiltonians from all layers. By contrast, as seen in Equation (16), the WNN method introduces multiplicative corrections to the affinity matrix elements, where these corrections for different layers are jointly computed from the local network connectivity information and macroscopic community structure of all layers, manifested as within- and cross-modality nearest neighbor prediction errors of individual layers’ clustering preferences.

Applying the SCML, Seurat WNN, and spectral clustering based on RNA or ADT alone to a single-cell CITE-seq dataset of cord blood mononuclear cells (Stoeckius *et al*., 2017) has shown that the SCML and Seurat WNN algorithms yield comparable clustering results, recapitulating the biologically meaningful separation of cell types and, at the same time, offering reconciliation of potential differences between the RNA and ADT modalities (Fig. 3, Table 1). Apparent differences between SCML and WNN results may inevitably arise because of the differences in their optimization functions and algorithmic details, but some of these differences may get resolved when resolution parameters and cluster numbers are adjusted. This result suggests that an ideal clustering of cells based on one modality may not always agree with that based on another modality and that an integrative clustering approach incorporating all available information better partitions the cells into biological cell types. Our result thus highlights the benefit of multi-omics research, as a single modality may be prone to noise or provide an incomplete description of single cells.

A limitation of the SCML algorithm is that it relies on the assumption that the majority of the modalities are informative of the true underlying clusters (Dong *et al*., 2013). If one of the modalities is very noisy, including this modality in the joint clustering may actually make it difficult to discover the true clusters detected by other modalities. The graph and embedding subspace obtained from the noisy modality may not capture the true cluster structure, but nevertheless will pull the consensus subspace towards the incorrect subspace on the Grassmannian manifold. Furthermore, if the clusters obtained from different modalities represent radically different partitions of the datasets, then finding a subspace between the individual embedding subspaces on the Grassmannian manifold may not represent a useful clustering of the dataset. This limitation is not, however, unique to the SCML algorithm and represents an assumption fundamental to many methods trying to integrate information across multiple modalities, as also noted as a limitation of the Seurat WNN algorithm (Hao *et al*., 2021).

By using a local modality weight, the Seurat WNN algorithm (Hao *et al*., 2021) may be able to handle differences in the informativeness of each modality for different cell types better than the SCML algorithm; however, while it may certainly be that cell-specific modality weights might aid in clustering certain cell types (Hao *et al*., 2021), we have shown that the SCML algorithm can uncover many of the same clusters without the need of cell-specific weights.

