## Supplementary Information for "Spectral clustering of single-cell multi-omics data on multilayer graphs"


#### S1 Supplementary Methods

##### S1.1 Hamiltonian from the random walk graph Laplacian

Graph Laplacians provide useful tools for signal processing on graphs. For example, given a function $\mathbf{f} : \mathcal{V} \to \mathbb{R}$ defined on the set $\mathcal{V}$ of $N$ nodes on a graph, we will show here that starting from the random walk graph Laplacian, $\mathbf{L}_{\text{rw}} \equiv \mathbf{D}^{-1}\mathbf{L} = \mathbf{I} - \mathbf{D}^{-1}\mathbf{A}$, we can construct a Hermitian operator whose expectation value with respect to $\mathbf{f}$ corresponds to the error of predicting the value of $\mathbf{f}$ at the nodes based on its values in the local neighborhood. Since the set of all functions on a finite set of $N$ elements is isomorphic to $\mathbb{R}^N$, we will use the convention $\mathbf{f} \in \mathbb{R}^N$ in our ensuing discussion.

The name random walk graph Laplacian stems from the fact that $\mathbf{P} \equiv \mathbf{D}^{-1}\mathbf{A}$ can be viewed as the transition matrix for a discrete-time homogeneous first-order Markov chain $X_t$, $t \in \mathbb{Z}_{\geq 0}$. That is, $\mathbf{P}$ is a stochastic matrix satisfying $\mathbf{P}\mathbb{1} = \mathbb{1}$,where $\mathbb{1} \equiv [1, \ldots, 1]^{\mathrm{T}}$; as a result, we have $\mathbf{L}_{\text{rw}}\mathbb{1} = 0$. The matrix elements of $\mathbf{P}$ satisfy,

$$P_{ij} \equiv P(X_{t+1} = j | X_t = i) = \frac{A_{ij}}{\mathrm{Deg}(v_i)}, \quad \pi_j(t+1) = \sum_i \pi_i(t)\, P_{ij},$$

where $\pi(t) = [\pi_1(t), \ldots, \pi_N(t)]$ is the row vector of probability of being at each indexed node at time $t$. The time evolution of $\pi(t)$ can be rewritten as

$$\pi(t+1) = \pi(t)\mathbf{P}, \quad \pi(t+1) - \pi(t) = \pi(t)(\mathbf{P} - \mathbf{I}) \equiv -\pi(t)\,\mathbf{L}_{\text{rw}}.$$

The stationary states $\pi_{\mathbf{s}}$ are left eigenvectors of $\mathbf{L}_{\text{rw}}$ with the (possibly degenerate) eigenvalue zero.

The action of $\mathbf{L}_{\text{rw}}$ on a vector-valued function $F \equiv (\mathbf{f}_1 \cdots \mathbf{f}_M) \in \mathbb{R}^{N \times M}$ on the graph is

$$(\mathbf{L}_{\text{rw}}F)_{im} = F_{im} - \sum_j \frac{A_{ij}}{\mathrm{Deg}(v_i)} F_{jm} \equiv (\mathbf{L}_{\text{rw}}\mathbf{f}_m)(v_i),$$

which is the difference between the true value of the vector on $v_i$, and the value "predicted" by the weighted sum of the function at the neighbors of $v_i$. The Locally Linear Embedding (LLE) algorithm

(Roweis and Saul, 2000) aims at minimizing the errors of the local neighborhood prediction, by minimizing the sum of their L2 norm squared,

$$\begin{aligned}\sum_i \|F(v_i) - \sum_j \frac{A_{ij}}{\mathrm{Deg}(v_i)} F(v_j)\|_2^2 &= \sum_i \|[\mathbf{L}_{\mathrm{rw}} F](v_i)\|_2^2 \\ =&\mathrm{tr}\left[F^{\mathrm{T}}\, \mathbf{L}_{\mathrm{rw}}^{\mathrm{T}} \mathbf{L}_{\mathrm{rw}} F\right] = \mathrm{tr}\left[F F^{\mathrm{T}}\, \mathbf{L}_{\mathrm{rw}}^{\mathrm{T}} \mathbf{L}_{\mathrm{rw}}\right].\end{aligned} \tag{1}$$

More generally, introducing another diagonal matrix, $\mathbf{W} = \mathrm{diag}\{w_i\}_{i=1}^N$, $w_i \in [0, 1], \forall i$, we have

$$\sum_i w_i^2 \|F(v_i) - \sum_j \frac{A_{ij}}{\mathrm{Deg}(v_i)} F(v_j)\|_2^2 = \mathrm{tr}\left[F^{\mathrm{T}}\, \mathbf{L}_{\mathrm{rw}}^{\mathrm{T}}\, \mathbf{W}^2\, \mathbf{L}_{\mathrm{rw}} F\right] = \mathrm{tr}\left[F F^{\mathrm{T}}\, \mathbf{L}_{\mathrm{rw}}^{\mathrm{T}}\, \mathbf{W}^2\, \mathbf{L}_{\mathrm{rw}}\right]. \tag{2}$$

In the language of quantum mechanics, we can think of Equation (2) as computing the expectation value of the Hamiltonian

$$\mathbf{H}_{\mathrm{LL}} = \mathbf{L}_{\mathrm{rw}}^{\mathrm{T}}\, \mathbf{W}^2\, \mathbf{L}_{\mathrm{rw}}, \tag{3}$$

with respect to a mixed state described by the density matrix $\rho = FF^{\mathrm{T}}/\mathrm{tr}[FF^{\mathrm{T}}]$ This intuition suggests that the WNN algorithm, designed to utilize the accuracy information about within-modality and cross-modality predictions can be understood as an optimization problem of searching for a constrained mixed-state density matrix minimizing the ensemble energy of a Hamiltonian operator closely related to the above $\mathbf{H}_{\mathrm{LL}}$.

#### S1.2 Truncated uniform density matrix and thermal density matrix

For a symmetric Hamiltonian operator $\mathbf{H}$ acting on an $N$-dimensional vector space, let $\lambda_m$ and $\mathbf{u}_m$, $m = 0, \ldots, N-1$, denote its eigenvalues, ordered as $\lambda_0 \leq \lambda_1 \leq \cdots \leq \lambda_{N-1}$, and corresponding orthonomal eigenvectors, respectively. A general density matrix is a Hermitian, positive semi-definite matrix with unit trace and can be expressed as

$$\rho = \sum_{m=0}^{N-1} p_m \mathbf{v}_m \mathbf{v}_m^{\dagger}, \quad p_m \geq 0,\ \forall m, \tag{4}$$

where $^{\dagger}$ indicates Hermitian conjugate, $\mathbf{v}_0, \ldots, \mathbf{v}_{N-1} \in \mathbb{C}^N$ are orthonormal vectors, and

$$\sum_{m=0}^{N-1} p_m = 1.$$

In the paper, we stated that the truncated mixed state of the lowest-frequency eigenvectors of $\mathbf{H}$ is the solution of the following trace minimization problem,

$$\min_{\rho} \mathrm{tr}\left[\rho\, \mathbf{H}\right], \quad \text{s.t.} \quad \|\rho\|_2 = \frac{1}{K}, \tag{5}$$

where $\|\rho\|_2 = \max_m\{p_m\}$ is the spectral norm of $\rho$, and $K$ is a fixed positive integer. This problem translates to

$$\min\left\{ \sum_{m=0}^{N-1} p_m \mathbf{v}_m^{\dagger}\, \mathbf{H}\, \mathbf{v}_m \;\middle|\; \mathbf{v}_n^{\dagger} \mathbf{v}_m = \delta_{nm}, \quad 0 \leq p_m \leq \frac{1}{K}, \forall m, \ \text{ and } \ \sum_{m=0}^{N-1} p_m = 1 \right\}.$$

Iterative search shows that the solution is

$$p_0 = p_1 = \cdots = p_{K-1} = \frac{1}{K}, \quad p_K = p_{K+1} = \cdots = p_{N-1} = 0, \quad \mathbf{v}_m = e^{i\theta}\, \mathbf{u}_m,$$

thus yielding the truncated uniform density matrix,

$$\boxed{\rho_K = \frac{1}{K}\sum_{m=0}^{K-1} \mathbf{u}_m \mathbf{u}_m^{\mathrm{T}}}. \tag{6}$$

The thermal mixed state is the solution to the trace minimization problem with a different constraint,

$$\min_{\rho} \operatorname{tr}[\rho\,\mathbf{H}], \quad \text{s.t.} \quad S_0 = -\operatorname{tr}[\rho \log \rho], \tag{7}$$

where $S_0 > 0$ is fixed. Defining the mean energy $E[\rho]$ and the von Neumann entropy $S[\rho]$ as

$$E[\rho] = \operatorname{tr}[\rho\,\mathbf{H}], \quad S[\rho] = -\operatorname{tr}[\rho \log \rho],$$

the above trace minimization problem is minimizing the energy while keeping the entropy fixed; it is also equivalent to maximizing the entropy while keeping the mean energy fixed,

$$\max_{\rho} \{-\operatorname{tr}[\rho \log \rho]\}, \quad \text{s.t.} \quad E_0 = \operatorname{tr}[\rho\,\mathbf{H}].$$

Introducing the Lagrange multiplier $\beta > 0$, the optimization problem is solved by finding the density matrix that minimizes the von Neumann free energy

$$F_\beta[\rho] = E[\rho] - \beta^{-1} S[\rho] = \operatorname{tr}[\rho\,\mathbf{H}] + \beta^{-1}\operatorname{tr}[\rho \log \rho] = \beta^{-1}\operatorname{tr}[\rho\,(\log \rho + \beta\,\mathbf{H})]$$

at some fixed $\beta$, which needs to be subsequently tuned to achieve the entropy $S_0$.

Define the density matrix $\rho_\beta$ as

$$\boxed{\rho_\beta \equiv \frac{1}{Z} e^{-\beta\,\mathbf{H}} = \frac{1}{Z}\sum_{m=0}^{N-1} e^{-\beta\lambda_m} \mathbf{u}_m \mathbf{u}_m^{\mathrm{T}}}, \quad \text{where } Z \equiv \operatorname{tr}\left[e^{-\beta\,\mathbf{H}}\right]. \tag{8}$$

This density matrix satisfies

$$\log \rho_\beta = -\beta\,\mathbf{H} - \log Z, \quad F_\beta[\rho_\beta] = -\beta^{-1}\log Z.$$

To show that it is the thermal equilibrium density matrix that minimizes the free energy at fixed $\beta$ (Preskill, 1998), we note that since the quantum relative entropy between any two density matrices $\rho$ and $\sigma$

$$S(\rho\|\sigma) = -\operatorname{tr}[\rho \log \sigma] - S[\rho] = \operatorname{tr}[\rho\,(\log \rho - \log \sigma)] \geq 0,$$

is always non-negative due to Klein's inequality, we must have

$$S(\rho\|\rho_\beta) = \operatorname{tr}[\rho\,(\log \rho + \beta\,\mathbf{H})] + \log Z = \beta\,(F_\beta[\rho] - F_\beta[\rho_\beta]) \geq 0.$$

Thus, the free energy attains its minimum when $\rho = \rho_\beta$ (Equation (8)). To find the value of $\beta$, we solve for $\beta$ in

$$S_0 = \beta \operatorname{tr}[\rho_\beta\,\mathbf{H}] + \log \operatorname{tr}\left[e^{-\beta\,\mathbf{H}}\right].$$

The thermal density matrix $\rho_\beta$ shares the eigenvectors with the original Hamiltonian, but gives higher probability weight to eigenstates with lower energy.

In an ideal case where the graph can be partitioned into $K$ subgraphs without cutting any existing edges, the graph Laplacian is block-diagonalizable with $K$ degenerate ground states. Note that at zero temperature, we have

$$\lim_{\beta\to\infty} e^{-\beta(\lambda_m-\lambda_0)} = \begin{cases} 1, & \lambda_m = \lambda_0 \\ 0, & \lambda_m > \lambda_0 \end{cases} \quad ;$$

that is, only the lowest energy eigenstates are possible. Thus if there exist $K$ degenerate ground states, then the thermal density matrix becomes

$$\lim_{\beta\to\infty} \rho_\beta = \frac{1}{K}\sum_{m=0}^{K-1} \mathbf{u}_m \mathbf{u}_m^{\mathrm{T}} = \rho_K. \tag{9}$$

In this case, the two choices of mixed states become identical. In reality, a Hamiltonian constructed from the graph Laplacian will have a unique ground state if the graph is connected. Experience shows that choosing finite but small temperature, so that $\beta \sim \sqrt{N} \gg 1$, produces a mixture dominated by several smallest eigenvalue eigenstates (see Supplementary Methods S1.5 for further discussion).

In our interpretation of the SCML algorithm and reformulation of the WNN method, we use a density matrix to represent the spectral embedding structure of each individual graph layer. The resulting set of density matrices then enters the optimization problem together with the single-layer Hamiltonian operators.

#### S1.3 Node-specific modality weights from minimizing the difference between within- and cross-modality prediction

In the paper, we reformulate the weighted nearest neighbor (WNN) analysis as an optimization problem in the Hamiltonian formalism of multilayer graphs, with the following measure of deviation between a pair $(s_1, s_2)$ of graph layers ,

$$\begin{aligned} &-\mathrm{tr}\left\{\left[\rho^{(s_1)} - \rho^{(s_2)}\right]\left[\mathbf{H}_{\mathrm{LL}}^{(s_1)} - \mathbf{H}_{\mathrm{LL}}^{(s_2)}\right]\right\} \\ =&\sum_{i=1}^{N}\left[w_i^{(s_1)}\right]^2\left\{\left[\Delta_i^{(s_2,s_1)}\right]^2 - \left[\Delta_i^{(s_1,s_1)}\right]^2\right\} + \sum_{i=1}^{N}\left[w_i^{(s_2)}\right]^2\left\{\left[\Delta_i^{(s_1,s_2)}\right]^2 - \left[\Delta_i^{(s_2,s_2)}\right]^2\right\}, \end{aligned}$$

where $\Delta_i^{(s_1,s_2)}$ is the cross-modality error of predicting on $s_1$ using the neighborhood information from $s_2$; the lower the value, the better we can use the neighboring nodes of $v_i$ in layer $s_2$ to predict its function value at $v_i$ in layer $s_1$. $\Delta_i^{(s_2,s_2)}$ is the baseline error from within-modality neighborhood prediction: the error of using the neighboring nodes of $v_i$ in layer $s_2$ to predict its function value at $v_i$ in the same layer $s_2$. The difference between the two terms informs how much weight should be assigned to the neighborhood structure of $v_i$ in layer $s_2$ when other layers are to be taken into account. That is, a small difference suggests that the connectivity structure of $v_i$ may be shared among multiple layers, and the weight of $v_i$ may be chosen to be relatively high in the consensus WNN; by contrast, a large difference suggests that the particular structure found in layer $s_2$ may be an outlier, and the weight of $v_i$ may need to adjusted down when constructing the WNN.

When $s_{\max} = 2$, we thus estimate the weight matrices by minimizing the above deviation measure, which amounts to solving the following optimization problem:

$$\begin{aligned}
\mathbf{W}_*^{(1)}, \mathbf{W}_*^{(2)} &= \underset{\mathbf{W}^{(1)}+\mathbf{W}^{(2)}=\mathbf{I}}{\arg\max} \ \operatorname{tr}\left\{\left[\rho^{(1)}-\rho^{(2)}\right]\left[\mathbf{H}_{\mathrm{LL}}^{(1)}-\mathbf{H}_{\mathrm{LL}}^{(2)}\right]\right\} \\
&= \underset{w_i^{(1)}+w_i^{(2)}=1,\forall i}{\arg\min} \ \sum_{i=1}^{N}\left[w_i^{(1)}\right]^2\left\{\left[\Delta_i^{(2,1)}\right]^2-\left[\Delta_i^{(1,1)}\right]^2\right\}+\sum_{i=1}^{N}\left[w_i^{(2)}\right]^2\left\{\left[\Delta_i^{(1,2)}\right]^2-\left[\Delta_i^{(2,2)}\right]^2\right\}.
\end{aligned} \tag{10}$$

Defining

$$D_i^{(s_1,s_2)} = \left[\Delta_i^{(s_1,s_2)}\right]^2 - \left[\Delta_i^{(s_2,s_2)}\right]^2,$$

the proposed solution is

$$w_{*i}^{(1)} = \begin{cases} \frac{1}{2} & \text{, if } D_i^{(1,2)} = D_i^{(2,1)} \\ \frac{D_i^{(1,2)}}{D_i^{(1,2)}+D_i^{(2,1)}} & \text{, if } D_i^{(1,2)}, D_i^{(2,1)} \geq 0 \\ \max\left\{\operatorname{sign}(D_i^{(1,2)} - D_i^{(2,1)}), 0\right\} & \text{, otherwise} \end{cases}$$

and $w_{*i}^{(2)} = 1 - w_{*i}^{(1)}$. Note that in the rare case when $D_i^{(1,2)}$ and $D_i^{(2,1)}$ are both negative and equal to each other, the solution to the optimization is not unique and could be either

$$(w_{*i}^{(1)}, w_{*i}^{(2)}) = (1,0) \ \text{ or } \ (w_{*i}^{(1)}, w_{*i}^{(2)}) = (0,1);$$

given the symmetry of the prediciton errors, we propose to use equal weights $w_{*i}^{(1)} = w_{*i}^{(2)} = 1/2$ in this case.

For the case of more than two layers, this scheme can be generalized by iterating over each pair of layers

$$\underset{\sum_{s=1}^{s_{\max}} \mathbf{W}^{(s)}=\mathbf{I}}{\arg\max} \sum_{(s_1,s_2)} \operatorname{tr}\left\{\left[\rho^{(s_1)}-\rho^{(s_2)}\right]\left[\mathbf{H}_{\mathrm{LL}}^{(s_1)}-\mathbf{H}_{\mathrm{LL}}^{(s_2)}\right]\right\},$$

and the solution is

$$w_{*i}^{(s)} = \begin{cases} \frac{1}{s_{\max}} & \text{, if } \sum_t D_i^{(t,s')} \text{ are equal for all } s' \\ \frac{\left(\sum_t D_i^{(t,s)}\right)^{-1}}{\sum_{s'}\left(\sum_t D_i^{(t,s')}\right)^{-1}} & \text{, if } \left(\sum_t D_i^{(t,s')}\right)^{-1} \geq 0 \ \forall s' \\ \frac{\mathbb{1}(\sum_t D_i^{(t,s')}=M_i)}{N_i} & \text{, otherwise} \end{cases} \tag{11}$$

where $\mathbb{1}$ is the indicator function, $M_i = \min_{s'}\left\{\sum_t D_i^{(t,s')}\right\}$, and $N_i$ is the number of elements in the set

$$\left\{s' \,\middle|\, \sum_t D_i^{(t,s')} = M_i\right\}.$$

#### S1.4 Application of our WNN reformulation to a synthetic data set of 3 layers

To demonstrate that our WLL reformulation of WNN can be applied to more than 2 graph layers, we tested our algorithm on a synthetic data set with known ground truth for the cluster labels. We used the function sklearn.datasets.make_blobs from the Python package scikit-learn, with options (n_samples=[50,30,20], n_features=2, cluster_std=2, shuffle=False, random_state=$n$), where $n$ is a different random state for each of the 3 runs; this approach generated, for each of the $s_{\max} = 3$ layers, $N = 100$ synthetic data points in two dimensions, forming 3 clusters of size 50, 30, and 20 with the cluster identity of a data point shared among the layers (Supplementary Figure 1A). The number $N = 100$ was chosen to make the individual data points visible in the heatmap representations of the graph Laplacian and the density matrix. We calculated the similarity graph for each layer. Since some clusters partially overlapped in each layer, separating the three clusters in individual layer was difficult. Follow the same framework of spectral analyses described above, we computed for each layer the graph Laplacian $\mathbf{L}^{(s)}$, the Hamiltonian $\mathbf{H}^{(s)}$, and the ensemble of Hamiltonian eigenstates $\rho = \sum_{m=0}^{N-1} p_m \mathbf{u}_m \mathbf{u}_m^{\mathrm{T}}$ (Supplementary Figure 1B,C,D); here, we chose the locally linear construction $\mathbf{H}_{\mathrm{LL}} \equiv [\mathbf{W}\mathbf{L}_{\mathrm{rw}}]^{\mathrm{T}} \mathbf{W}\mathbf{L}_{\mathrm{rw}}$ with $\mathbf{W} = \mathbf{I}$, and the thermal mixture with $p_m = e^{-\beta\lambda_m} \Big/ \sum_{m'=0}^{N-1} e^{-\beta\lambda_{m'}}$, using $\beta = \sqrt{N} = 10$, as we have suggested. We then calculated the node-specific weights for each layer using Equation (11), constructed the combined Hamiltonian $\mathbf{H}_{\mathrm{WLL}}$ for the multilayer graph, and performed $K$-means clustering in the $K = 3$ dimensional low-frequency eigenspace of $\mathbf{H}_{\mathrm{WLL}}$. The normalized mutual information (NMI) between the true cluster labels and the clustering labels obtained by our method was 0.914. The multilayer density matrix $\rho$ and feature map $\Phi$ (after dimension reduction using UMAP) at different values of $\beta$ are shown in Supplementary Figure 2.

#### S1.5 Effects of the inverse temperature $\beta$ on the reformulated WNN algorithm

As previously mentioned, we suggest using the inverse temperature $\beta = \sqrt{N}$, where $N$ is the number of single cells, when constructing the thermal ensemble of Hamiltonian eigenstates. Here, we discuss the effect of varying $\beta$ on spectral clustering results.

As described in Section S1.2, the thermal density matrix $\rho$ shares the eigenvectors with the original Hamiltonian, but gives higher weight to states with a lower eigenvalue. In the limit $\beta \to 0$,

$$p_m = \frac{e^{-\beta\lambda_m}}{\sum_{m'=0}^{N-1} e^{-\beta\lambda_{m'}}} = \frac{1}{N}, \quad \rho = \frac{1}{N}\sum_{m=0}^{N-1} \mathbf{u}_m \mathbf{u}_m^{\mathrm{T}},$$

and all states become equiprobable; in the opposite limit $\beta \to \infty$, if there exist $K$ degenerate ground states (i.e., the lowest eigenvalue $\lambda_0$ has multiplicity $K$), we have

$$p_m = \frac{\mathbb{1}(\lambda_m = \lambda_0)}{K}, \quad \rho = \frac{1}{K}\sum_{m=0}^{K-1} \mathbf{u}_m \mathbf{u}_m^{\mathrm{T}},$$

suppressing all higher eigenvalue states. In practice, the Hamiltonian constructed from the graph Laplacian will not have degenerate ground states if the graph has a single connected component, but it may have several low eigenvalues that are approximately degenerate.

The spectrum of the symmetric graph Laplacian $\mathbf{H}_{\mathrm{SC}} = \mathbf{D}^{-1/2}\,\mathbf{L}\,\mathbf{D}^{-1/2}$ is known to approximate the normalized cut minimization problem on a graph. Assuming that the number $K$ of clusters satisfies the condition $K \ll N$, the total weight of inter-cluster links would be of order $O(1)$, while the total weight of intra-cluster links would also be of order $O(N)$. Since solving $\mathbf{H}_{\mathrm{SC}}$ approximately solves the normalized cut minimization problem, the first few lowest eigenvalues thus satisfy $\lambda_m \sim O(1/N)$, $m = 0, \ldots, K-1$, while the higher eigenvalues are expected to behave as $\lambda_m \sim O(1)$, $m \geq K$. For too small a value of $\beta$, all modes become equiprobable, and the contrast between low-frequency grouping of nodes into clusters and high-frequency noise of individual nodes is lost, potentially leading to the blurring of overall community structure. In the opposite case of too large a value of $\beta$, only the smallest eigenvalue would survive, potentially leading to the loss of subcluster structure within larger clusters and yielding only a low-resolution clustering result. In order to detect the proper macroscopic clustering structure of a graph, we thus need $\beta \ll N$ to retain the first few low-frequency modes, while we also need to impose the condition $\beta \gg 1$ to suppress high-frequency noisy modes.

We found that the suggested choice of $\beta = \sqrt{N}$ performs well in detecting clusters. We demonstrated the effect of varying $\beta$ using the synthetic triple-layer data set of $N = 100$ points described above (Supplementary Figure 2). The normalized mutual information (NMI) between the true cluster labels (indicated by different colors in the UMAP plots) and the clustering labels obtained by our method were 0.886, 0.914, and 0.870, respectively, for $\beta = 1, 10, 100$. From the feature maps and the multilayer density matrix, we observed that a small value of $\beta$ resulted in low contrast between the clusters, while a large value of $\beta$ resulted in loss of resolution in the density matrix. By contrast, the recommended value of $\beta = \sqrt{N} = 10$ performed the best in this simulation study. We performed a similar analysis for a subset of $N = 100$ cells from the CBMC data set (Stoeckius *et al.*, 2017), under a wide range of inverse temperature values $\beta = 0.1, 1, 10, 100, 1000$ (Supplementary Figure 3), and found that the recommended value of $\beta = 10$ again performed well in terms detecting broad clusters and also retaining the resolution of subclusters.

#### S1.6 Effects of the hyperparameter $\alpha$ on SCML clustering results

Given a set of graph layer adjacency matrices, the SCML algorithm requires the choice of one hyperparameter, $\alpha \in \mathbb{R}_{\geq 0}$, that sets the balance between the summed graph Laplacians and the projection distance to the individual graph layer embeddings (Dong *et al.*, 2013). A graph adjacency matrix encodes information about the microscopic connectivity of nodes, while a spectral embedding subspace encodes information about grouping nodes into macroscopic communities. The $\alpha$ parameter controls the compromise between these two representations of multilayer graphs. As noted in (Dong *et al.*, 2013), choosing a small value of $\alpha$ amounts to averaging the adjacency matrices of multilayer graphs, while choosing a large value of $\alpha$ amounts to averaging the spectral kernel (Gram matrix of spectral embedding subspace matrices) and using the resulting average as the consensus adjacency matrix. An intermediate value of $\alpha$ retains information about the layer-specific local connectivity of nodes, while also forcing the algorithm to find a spectral embedding subspace that is close to all individual subspaces of the layers.

To evaluate the effect of the hyperparameter in clustering the CBMC CITE-seq data set (Stoeckius *et al.*, 2017), we performed a grid search over a range of values for the hyperparameter $\alpha$ and calculated the silhouette score for the RNA, ADT, and embedding distance (Supplementary Figure 4A,B). Within the grid search range, we observed regions of values of the hyperparameter $\alpha$ within which the clustering results were similar (Supplementary Figure 4C,D). This analysis suggested there were multiple distinct joint embeddings corresponding to different values of $\alpha$, but each of these embeddings exhibited a range of $\alpha$ values that gave nearly identical clustering results. As such, a given clustering result must be chosen, but there was a range of values for which this clustering result was not sensitive to the value of $\alpha$.

We observed the highest silhouette score for RNA was obtained when $\alpha = 0.1$ and for ADT when $\alpha = 7.5$ (Supplementary Figure 4A). For both of these values, however, the silhouette score for the other modality was much lower than that obtained for other values of the hyperparameter $\alpha$. A large value of $\alpha$ ($\alpha \geq 100$) resulted in an ADT silhouette score near the optimal value while also obtaining a much higher RNA silhouette score compared to the optimal $\alpha$ value for ADT. A large value of $\alpha$ corresponded to a higher emphasis on minimizing the projection distance between the joint SCML embedding and the spectral embeddings obtained from the RNA and ADT graph layers. This suggested balancing the information from the individual graph layers provided better separation of the clusters than obtaining an optimal embedding based on the summed graph Laplacians. While other values of the hyperparameter $\alpha$ resulted in lower silhouette scores, some of the alternate clustering results were still biologically motivated. For example, the clustering result using a value of $\alpha = 1$ did not partition the NK cells into two subtypes, but instead included a separate cluster for pDCs (Seurat cluster 10) (Supplementary Figure 5). For a general data set, we recommend user exploration based on prior biological knowledge to choose the hyperparameter $\alpha$.

##### S1.7 Application to human bone marrow mononuclear cells (BMNC) CITE-seq data set

In addition to the CBMC dataset, we applied spectral clustering based on RNA, ADT, and the SCML algorithm to cluster cells in the BMNC data set from (Stuart *et al.*, 2019). The BMNC data set was used in (Hao *et al.*, 2021) to benchmark the WNN graph construction and clustering. We reproduced the Seurat WNN results for this data set by following the Seurat Weighted Nearest Neighbor vignette. The dataset was downloaded as a Seurat object using the SeuratData package. Following the vignette, we used a resolution of 2.0 for clustering, which resulted in 40 clusters.

To apply the spectral clustering methods to this data set, the RNA and protein counts for the filtered Seurat data set were saved from the downloaded Seurat object. The same processing and graph construction used for performing spectral clustering of the CBMC dataset were applied to construct the RNA and protein surface marker graphs, with the exception that 50 principal components were used for the RNA feature profiles. The SCML clustering was performed following the same protocol as for the CBMC dataset with a hyperparameter of $\alpha = 0.1$. For each method, the cells were clustered into 40 clusters. Clustering results for each of the methods are shown projected onto the Seurat WNN UMAP projections in Supplementary Figure 8.

The clustering results for the BMNC data set were difficult to compare via simple visual inspection, as the cluster colors were too similar for a large number of clusters. To compare the clustering results in a quantitative way, we used the normalized mutual information (NMI) score (`sklearn.metrics.normalized_mutual_info_score`) to measure the pairwise agreement between clustering methods (Supplementary Table 2). The NMI scores demonstrated that both the Seurat WNN and SCML clustering results had greater agreement with both modalities than clustering based on the alternative modality. Additionally, the NMI score between the Seurat WNN and SCML algorithms showed that these clustering methods agreed the best among all pairs of methods compared.

In Hao *et al.* (2021) the correlation between true protein ADT features and predicted profiles obtained by averaging over nearest neighbor profiles in the WNN graph was used to evaluate the WNN performance. To compare the spectral clustering methods using this same approach, we used the adjacency matrices for a single modality or a modified adjacency matrix obtained from the modified SCML graph Laplacian to predict the feature profiles. For a single modality, the ADT profiles were predicted as:

$$F_{\text{pred}} = \mathbf{D}^{-1}\mathbf{A}F, \tag{12}$$

where $\mathbf{A}$ is the adjacency matrix for the corresponding modality, $\mathbf{D}$ the corresponding degree matrix, and $F$ the matrix of CLR transformed ADT profiles with rows corresponding to cells and columns to different surface proteins. For the SCML algorithm, a joint adjacency matrix was obtained from the SCML-modified graph Laplacian:

$$\mathbf{A}_{\text{SCML}} = \text{diag}\left(\sum_{i=1}^{N}(\mathbf{L}_{\text{SCML}})_{i,1}, \cdots \sum_{i=1}^{N}(\mathbf{L}_{\text{SCML}})_{i,N}\right) - \mathbf{L}_{\text{SCML}}. \tag{13}$$

The predicted protein features were then obtained using Equation 12 with the SCML joint adjacency matrix (Equation 13). The correlation across cells between the predicted features (columns of $F_{\text{pred}}$) and the true features (columns of $F$) was taken for each of the RNA, ADT, and SCML adjacency matrices (Supplementary Figure 9). Feature prediction for the WNN algorithm was obtained by averaging the features over the 20 nearest neighbors on the WNN graph, as described in (Hao *et al.*, 2021). Pearson and Spearman correlation coefficients were obtained using scipy.stats.pearsonr and scipy.stats.spearmanr with default values, respectively. The correlation between predicted and true features for the WNN algorithm was comparable to that for the SCML method (Supplementary Figure 9).

### S2 Supplementary Figures

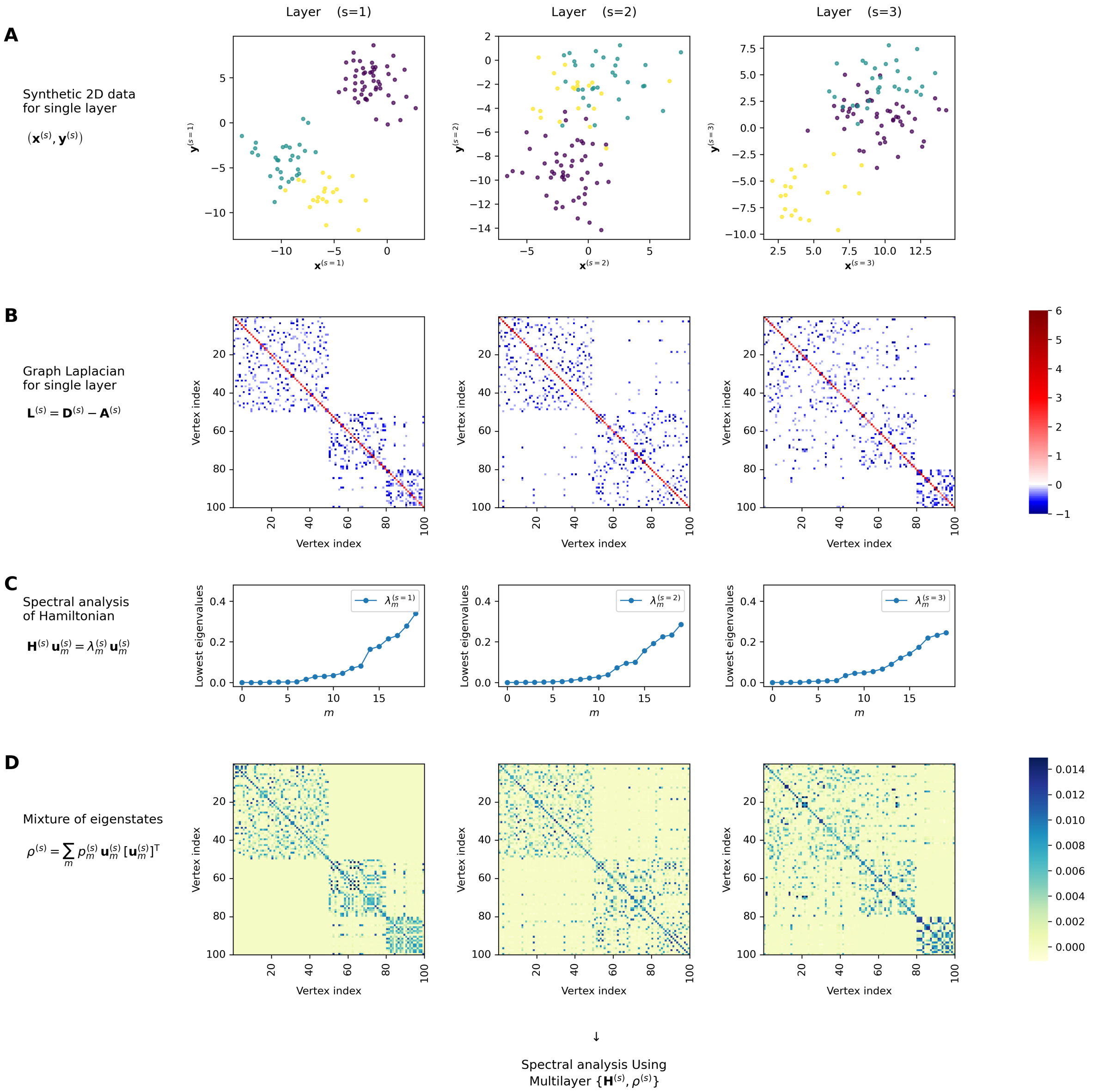


**Supplementary Figure 1:** Illustration of our reformulation of the WNN method applied to a triple-layer graph. For each layer, $N = 100$ synthetic data points in two dimension were randomly generated, forming 3 clusters of size 50, 30, and 20, respectively. (A) Heatmaps displaying the graph Laplacian $\mathbf{L}^{(s)}$ constructed from the pairwise similarity of the data in each layer. (B) The spectrum of the Hamiltonian $\mathbf{H}^{(s)}$ constructed from the graph Laplacian of each layer. The locally linear construction $\mathbf{H}_{\mathrm{LL}} \equiv [\mathbf{W}\mathbf{L}_{\mathrm{rw}}]^{\mathrm{T}}\mathbf{W}\mathbf{L}_{\mathrm{rw}}$ with $\mathbf{W} = \mathbf{I}$ was chosen here. (C) Heatmaps showing the density matrix representation of a mixed Hamiltonian eigenstates for each layer. The thermal mixture with $p_m = e^{-\beta\lambda_m} \Big/ \sum_{m'=0}^{N-1} e^{-\beta\lambda_{m'}}$ was chosen here.

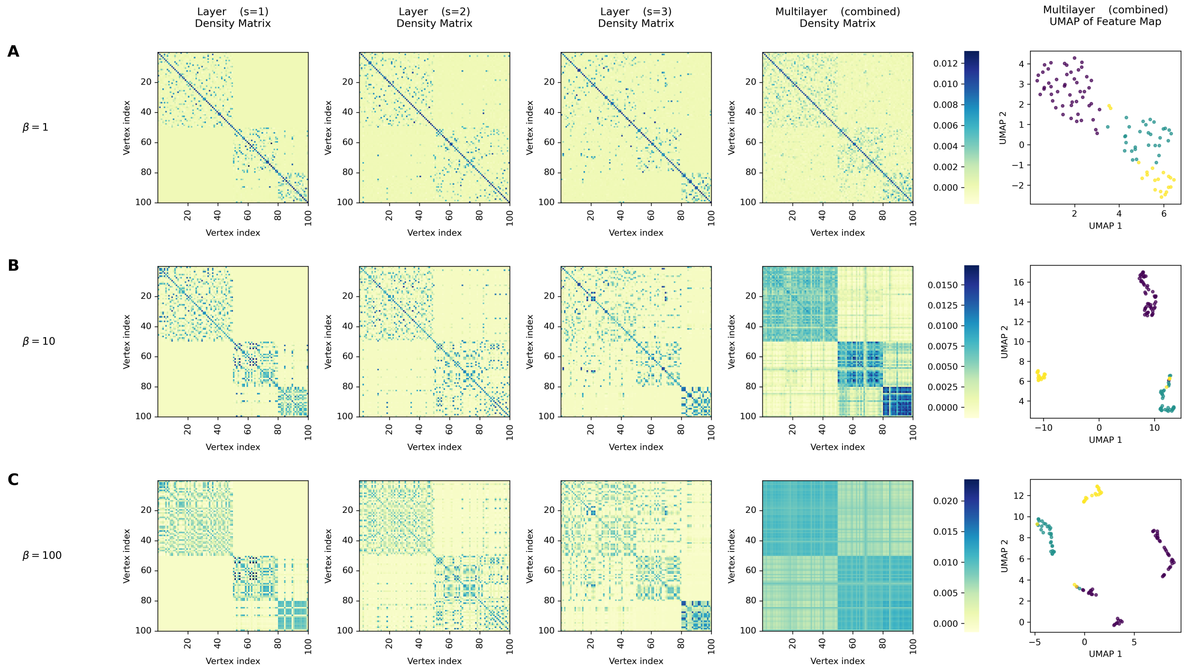


**Supplementary Figure 2:** Effects of varying the inverse temperature $\beta$ on the simulated data from Supplementary Figure 1. Density matrices representing the probability mixtures of the Hamiltonian eigenstates $\rho = \sum_{m=0}^{N-1} p_m \mathbf{u}_m \mathbf{u}_m^{\mathrm{T}}$ at values (A) $\beta = 1$, (B) $\beta = 10$, and (C) $\beta = 100$ are shown for each layer and the combined multilayer graph. The last column shows the UMAP projection of the corresponding feature map, $\Phi(v_i) = (\sqrt{p_0}\mathbf{u}_0(v_i), \ldots, \sqrt{p_{N-1}}\mathbf{u}_{N-1}(v_i))$, for the multilayer graph. The colors indicate the true cluster identities. The normalized mutual information (NMI) between the true cluster labels and the clustering labels obtained by our method are 0.886, 0.914, and 0.870 for $\beta = 1, 10$, and 100, respectively, supporting the validity of using the recommended value of $\beta \sim \sqrt{N} = 10$.

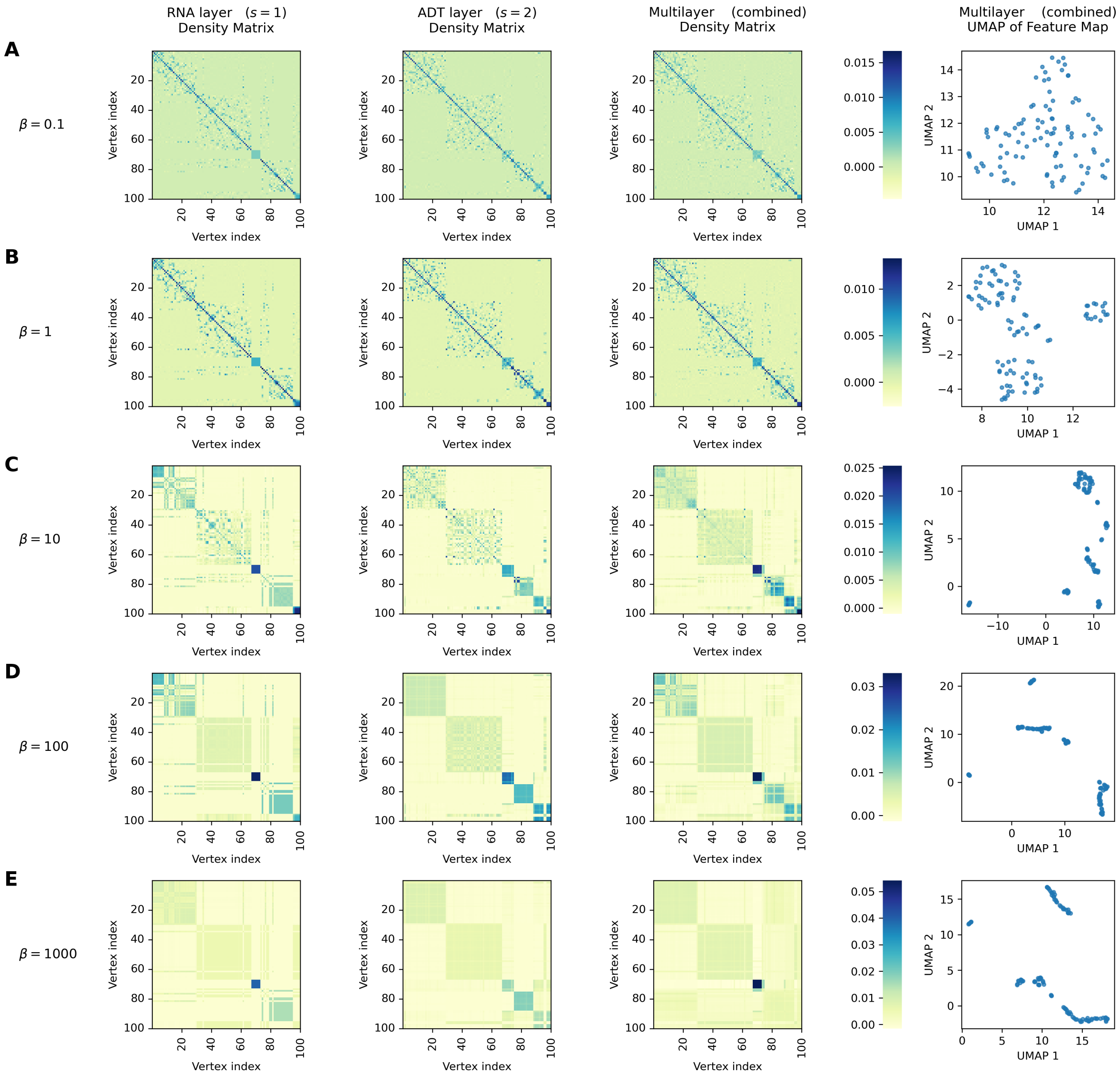


**Supplementary Figure 3:** Effects of varying the inverse temperature $\beta$ on the human CBMC data. Our reformulation of WNN using the thermal ensemble $p_m = e^{-\beta\lambda_m} \Big/ \sum_{m'=0}^{N-1} e^{-\beta\lambda_{m'}}$ was applied to a subset of $N = 100$ cells randomly selected from the CBMC CITE-seq data (Stoeckius *et al.*, 2017). Density matrices representing the probability mixtures of the Hamiltonian eigenstates $\rho = \sum_{m=0}^{N-1} p_m \mathbf{u}_m \mathbf{u}_m^{\mathrm{T}}$ at values (A) $\beta = 0.1$, (B) $\beta = 1$, (C) $\beta = 10$, (D) $\beta = 100$, and (E) $\beta = 1000$, are shown for each layer and the combined multilayer graph. The last column shows the UMAP projection of the corresponding feature map, $\Phi(v_i) = (\sqrt{p_0}\mathbf{u}_0(v_i), \ldots, \sqrt{p_{N-1}}\mathbf{u}_{N-1}(v_i))$, for the multilayer graph. The recommended value of $\beta \sim \sqrt{N} = 10$ is seen to detect broad clusters and also retain the resolution of subclusters.

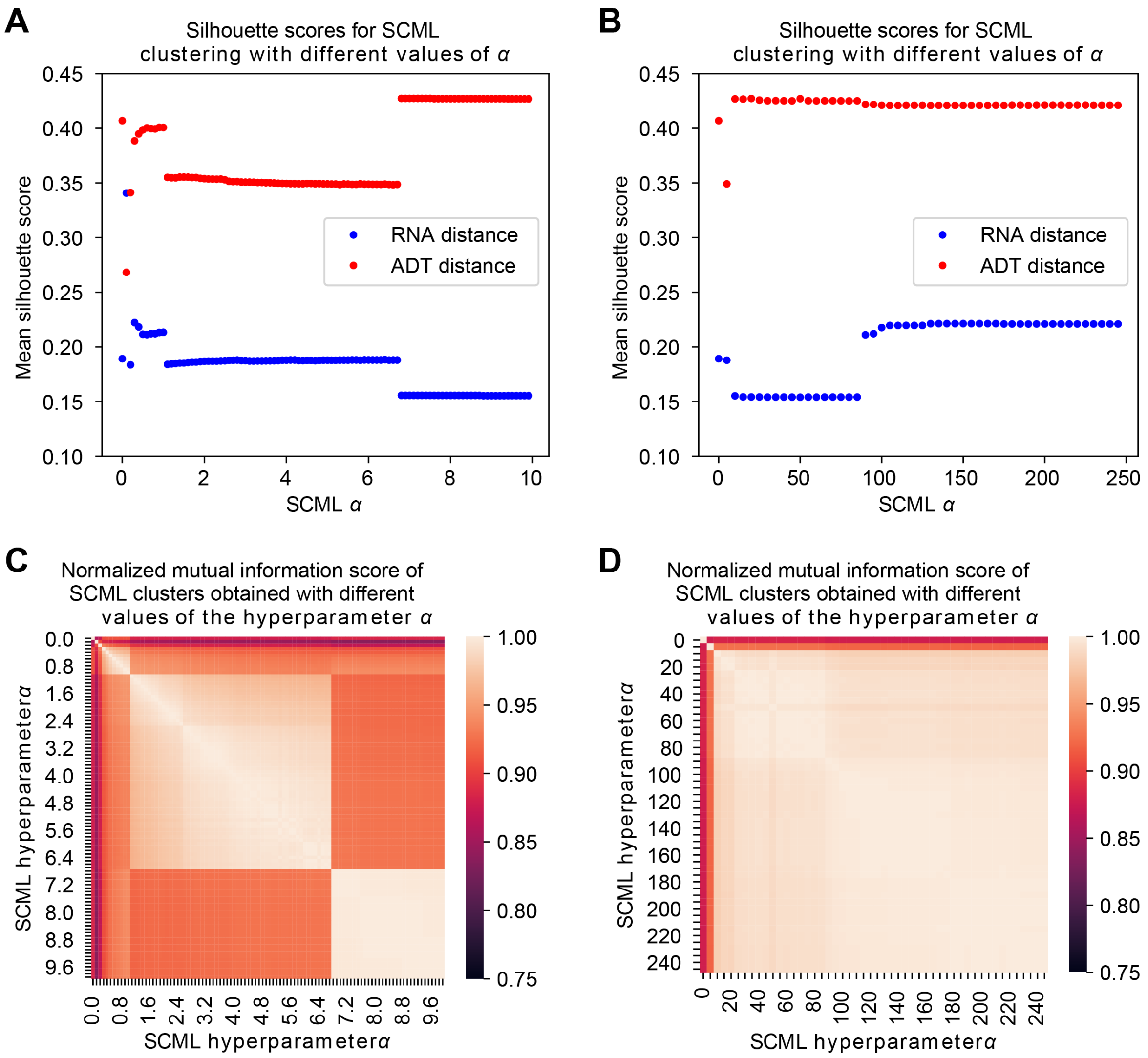


**Supplementary Figure 4:** SCML hyperparameter $\alpha$ search for the CBMC dataset (Stoeckius *et al.*, 2017). (**A**) Silhouette scores calculated using the pairwise ADT CLR distance matrix, RNA PCA distance matrix, and embedding distance matrix and clustering results obtained from the SCML algorithm with different values of the hyperparameter $\alpha$. 100 values of $\alpha$ ranging from 0 to 10 with interval 0.1. (**B**) Same as panel (A) but for values of $\alpha$ ranging from 0 to 250 with a step size of 5. (**C**) Pairwise normalized mutual information (NMI) score for SCML clusteing results obtained using different values of $\alpha$. The sample $\alpha$ values are the same used in panel (A). (**D**) The same as panel (C), but for the $\alpha$ values used in panel (B).

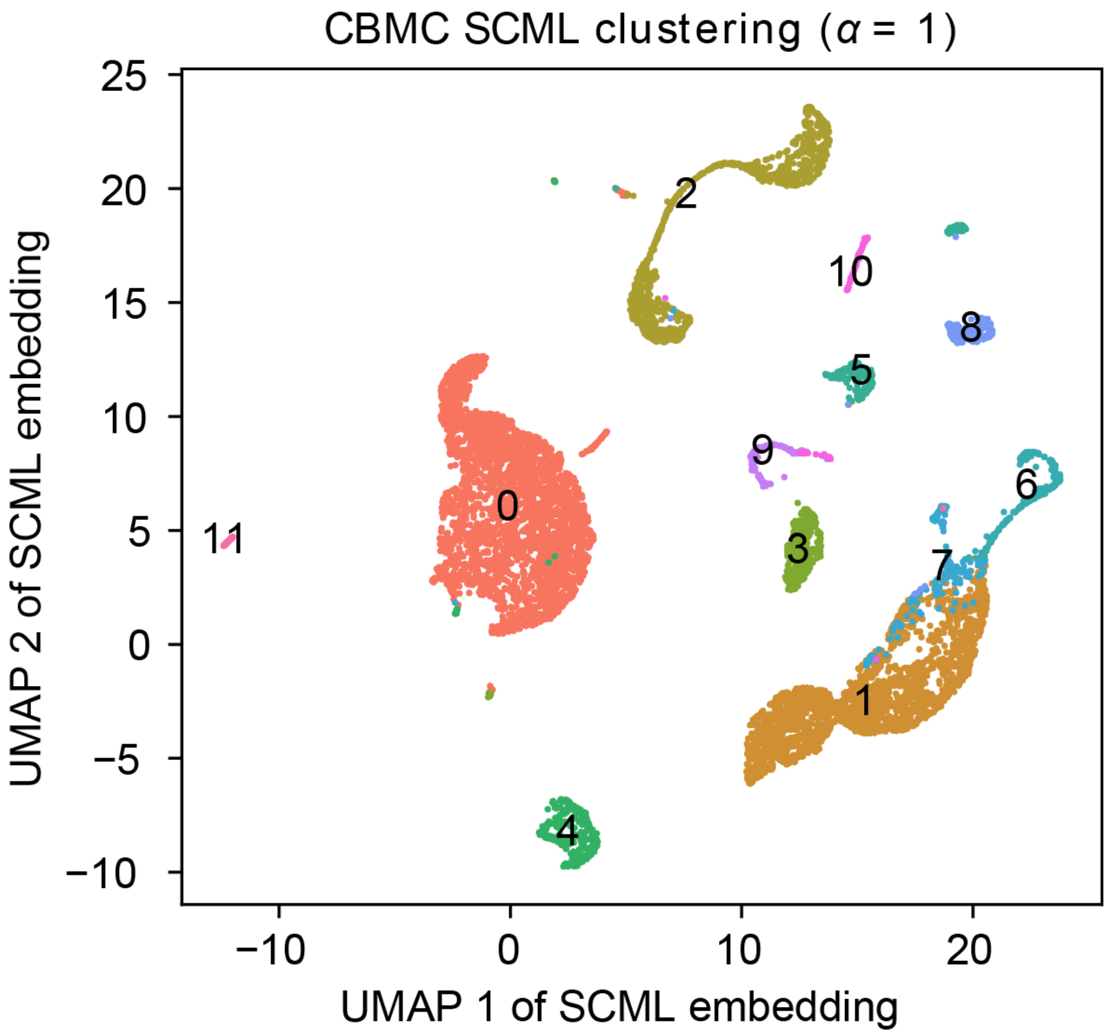


**Supplementary Figure 5:** SCML clustering of the CBMC dataset using a hyperparameter of $\alpha = 1$.

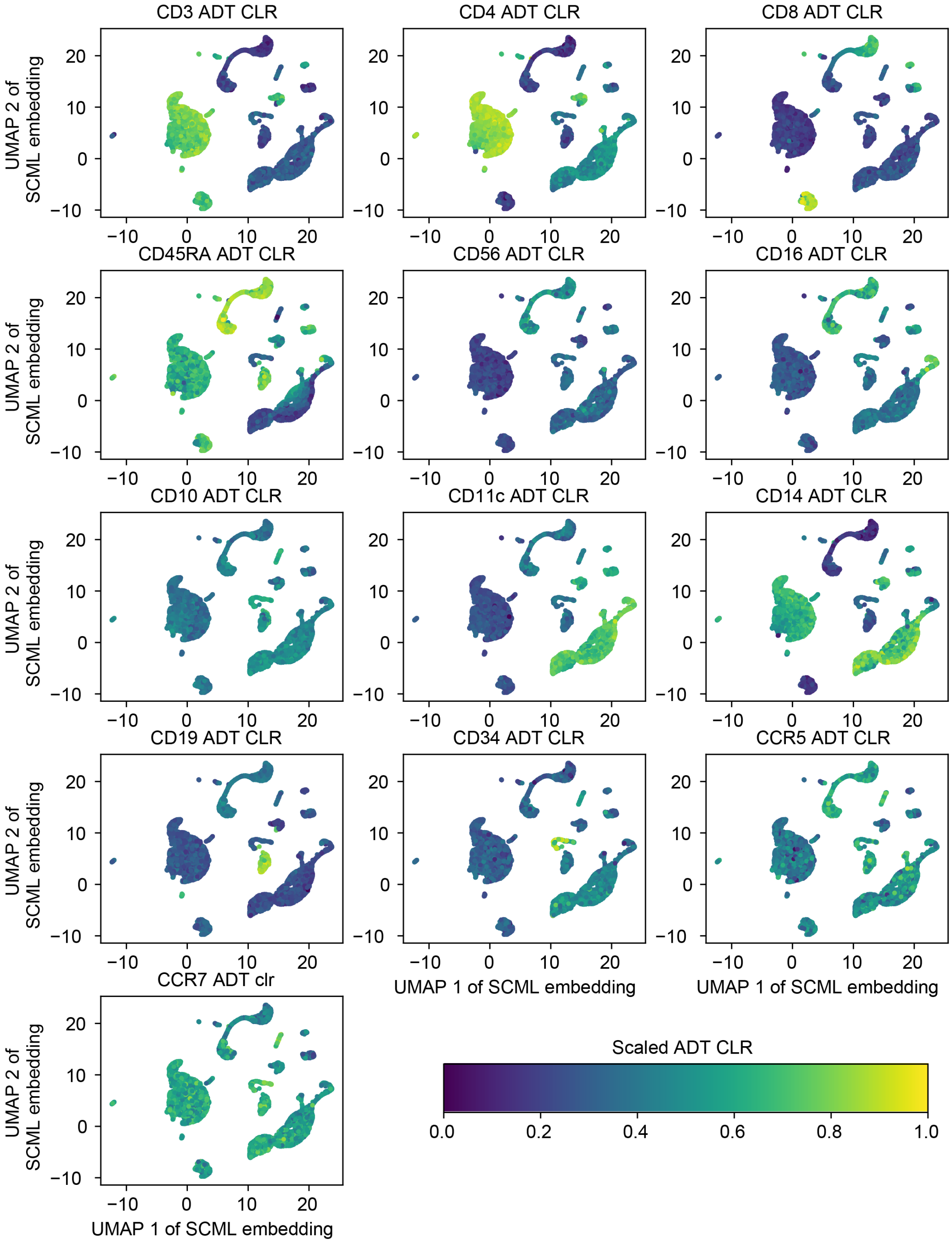


**Supplementary Figure 6:** Distribution of ADT CLR transformed profiles projected onto the SCML UMAP. For each of the 13 antibodies, the range of values was shifted and rescaled to fall within the range [0,1].

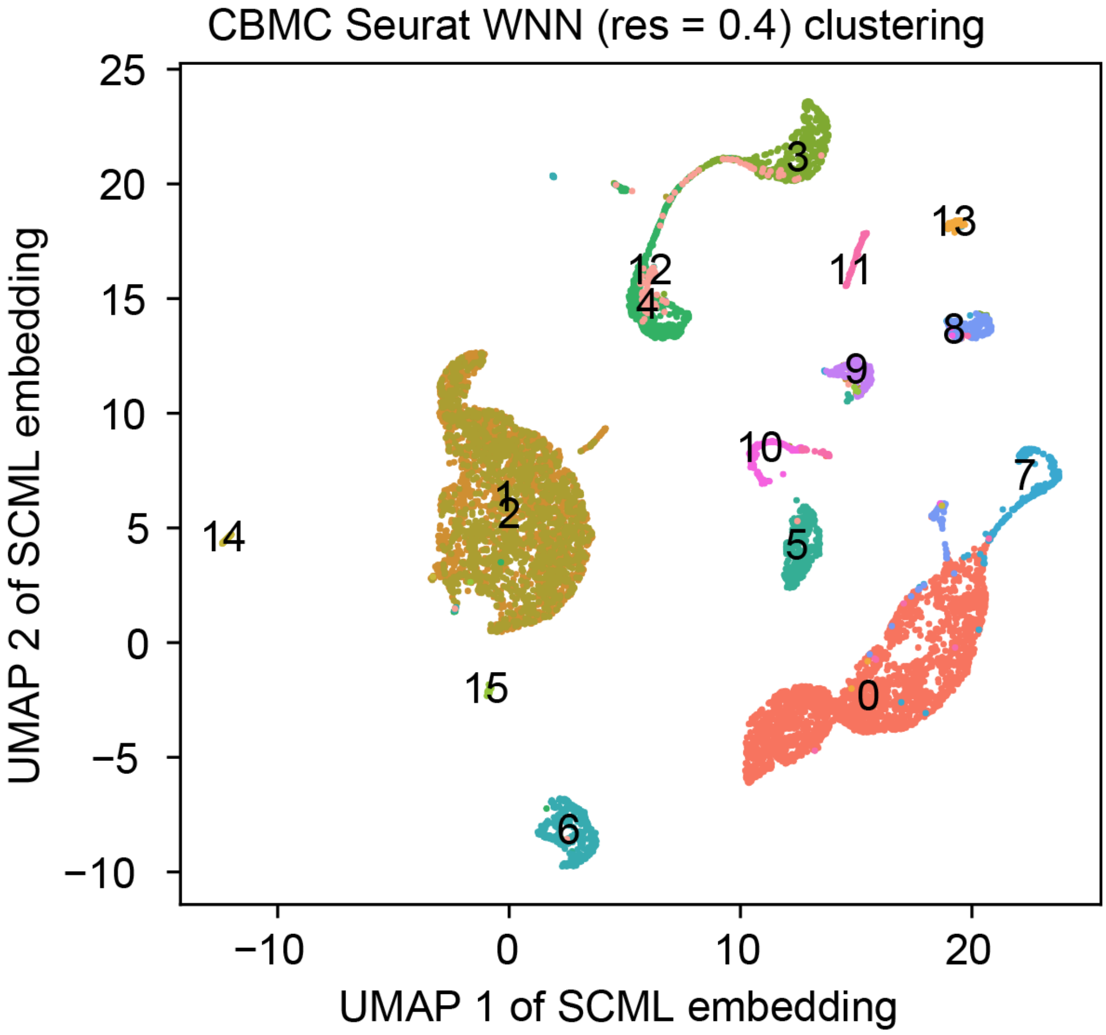


**Supplementary Figure 7:** Seurat WNN clustering with resolution 0.4. The clustering results are projected onto the two dimensiona UMAP projection obtained from the SCML embedding.

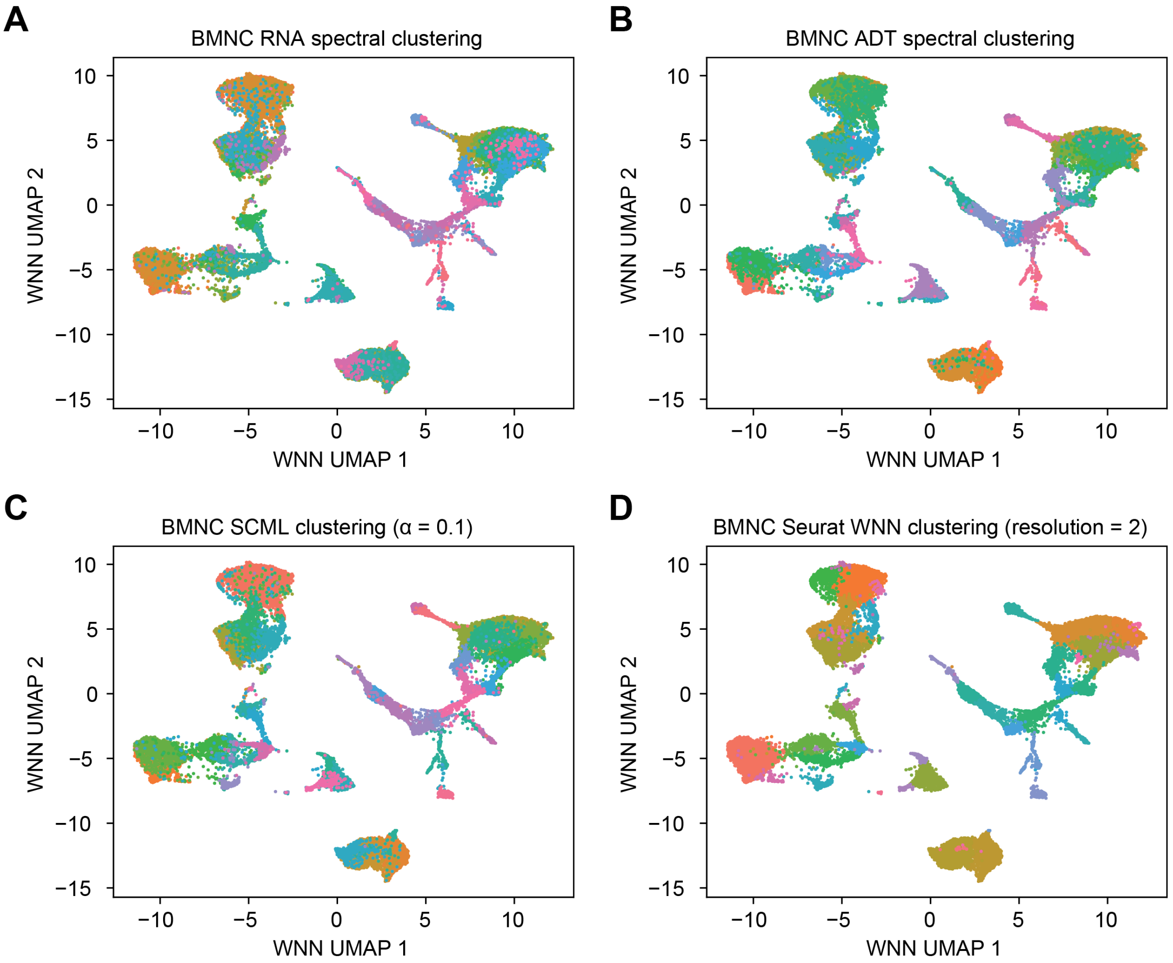


**Supplementary Figure 8:** Clustering results for the BMNC dataset (Stuart *et al.*, 2019) projected onto Seurat WNN UMAP. (**A**) RNA spectral clustering into 40 clusters. (**B**) ADT spectral clustering into 40 clusters. (**C**) SCML clustering into 40 clusters. (**D**) Seurat WNN clustering using a resolution of 2, which clustered the cells into 40 clusters.

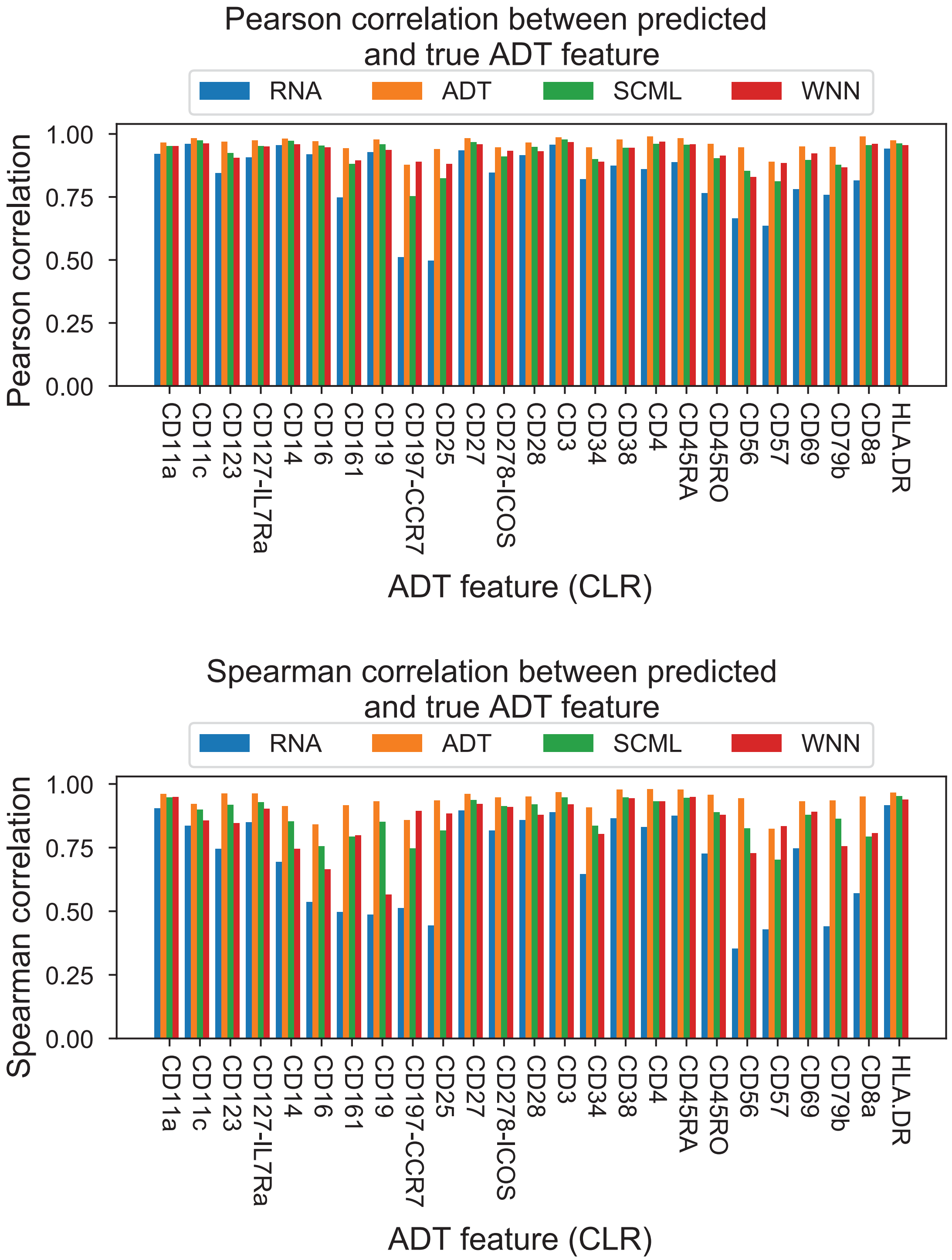


**Supplementary Figure 9:** Correlation between the predicted and true ADT features for BMNC dataset (Stuart *et al.*, 2019). (**A**) The Pearson correlation coefficient between the predicted and true ADT features for the BMNC dataset. Predictions were obtained for each of the RNA adjacency matrix, ADT adjacency matrix, SCML joint adjacency matrix, and based on Seurat WNN nearest neighbors. ADT features are the CLR transformed ADT features used for graph construction. (**B**) The same as (**A**) but showing the Spearman correlation between the predicted and true features.

#### S3 Supplementary Table

**Supplementary Table 1:** CBMC silhouette scores for different clustering methods and distance measures

| Clustering method | RNA distance | ADT distance |
|---|---|---|
| RNA spectral | 0.3696 | 0.2496 |
| ADT spectral | 0.08461 | 0.3941 |
| SCML ($\alpha = 100$) | 0.2177 | 0.4211 |
| SCML ($\alpha = 100$) (merged T cells) | 0.3302 | 0.3486 |
| Seurat WNN (res=0.2) | 0.2406 | 0.3977 |
| Seurat WNN (res=0.2) (merged T cells) | 0.2788 | 0.2998 |

**Supplementary Table 2:** BMNC normalized mutual information scores between clustering results obtained with different methods

| | RNA | ADT | SCML | WNN |
|---|---|---|---|---|
| RNA | 1.0 | 0.567 | 0.667 | 0.629 |
| ADT | 0.567 | 1.0 | 0.744 | 0.751 |
| SCML | 0.667 | 0.744 | 1.0 | 0.765 |
| WNN | 0.629 | 0.751 | 0.765 | 1.0 |
